# Development of an Integrated Single-Cell and Spatial Transcriptomics Atlas of Healthy Human Skin Focusing on the Pilosebaceous Unit

**DOI:** 10.1101/2025.09.09.675235

**Authors:** Tolga Düz, Daniel Torocsik, Sonia M. E. Momnougui, Mhaned Oubounyt, Benjamin Al, Stefan Gallinat, Jan Baumbach, Nicholas Holzscheck

**Author notes:** Corresponding author: Tolga Düz, Beiersdorfstraße 1-9, 20245 Hamburg, Germany;.

## Abstract

Single-cell and spatial transcriptomics have transformed our ability to chart human tissue organization at unprecedented resolution. These technologies enable the construction of high-quality reference atlases, essential for mapping healthy tissue architecture and identifying robust gene markers. We developed the healthy Human Skin Cell Atlas (HSCA), systematically integrating 34 public datasets and totaling 821,464 cells, with curated metadata and harmonized cell type nomenclature to ensure consistency. We place particular emphasis on the pilosebaceous unit, a key epithelial structure critical for both homeostasis and pathology. While prior studies captured the interfollicular epidermis and immune landscape in detail, deeper hair follicle regions remained under-characterized. By leveraging high-resolution spatial transcriptomics (Visium HD), we spatially resolved and transcriptionally defined the lower hair follicle compartments and pinpointed signalling hubs. Furthermore, the HSCA enables the detection of cell types not visible in standalone datasets, such as Merkel cells. Our results illustrate the value of the integrated single-cell atlas and spatial data in refining tissue organization and highlight the PSU as a complex and diverse epithelial niche.

## 1 INTRODUCTION

The human skin is the body’s largest organ and plays essential roles in barrier protection, immune defense, and sensory perception^1^. Its structural complexity is reflected in diverse compartments, including the pilosebaceous unit (PSU)^2^, which represents a key epithelial appendage involved in homeostasis, regeneration, and skin diseases^3^.

Breakthroughs in single-cell^4,5^ and spatial transcriptomics^6^ have made it possible to dissect the cellular and molecular composition of tissues at unprecedented resolution. These technologies have enabled the creation of reference atlases^7^, which serve as foundational resources for biomarker discovery, patient stratification, and disease modeling^8^. For such atlases to be effective, they must integrate data across donors, anatomical sites, and measurement modalities, while controlling for technical batch effects and preserving genuine biological variation^9^.

Community-driven efforts, including the Human Cell Atlas, have already delivered integrated, organ-scale atlases for several tissues, as summarized by Hrovatin, et al.^9^. A prominent example is the groundbreaking Human Lung Cell Atlas^10^, which provided a valuable framework for constructing integrated atlases. In the context of skin, a roadmap for a Human Skin Cell Atlas has been proposed^11^, reflecting the growing demand for a comprehensive skin reference. A platform has also been developed to provide curated single-cell RNA sequencing (scRNA-seq) datasets for skin^12^. Nevertheless, a full-thickness, integrated atlas of healthy human skin is still missing.

While previous studies have characterized specific features of human skin^13–16^, they often focus on isolated anatomical regions and rely on limited sampling strategies and donor diversity. Some large-scale datasets have emerged, including those with diseased samples^17^ or spanning multiple anatomical sites^18^, but these likewise represent isolated resources rather than unified, integrated atlases.

There are integrative approaches that have begun to address these limitations. For instance, Thrane, et al. ^19^ demonstrated the potential of integrative analyses for resolving the PSU, but were limited to four scRNA-seq datasets and insufficient spatial resolution. A fibroblast atlas^20^ leveraged extensive data to reveal five major fibroblast axes, yet focused solely on fibroblasts and did not provide a comprehensive atlas of all skin cell types or their spatial organization.

Importantly, none of these studies performed a systematic, quantitative validation of integration performance. As single-cell datasets expand in size and heterogeneity, robust, benchmarked integration is essential to retain genuine biological variation while correcting batch effects^9^. The scIB pipeline offers a rigorous framework to evaluate integration across algorithms and to balance correction against biological fidelity^21^.

Here, we present a fully integrated scRNA-seq atlas of healthy human skin across multiple anatomical sites (Fig. 1). By harmonizing 34 scRNA-seq datasets from 160 donors we generated the Human Skin Cell Atlas (HSCA) comprising 821,464 cells and 110 distinct cell types. This dataset is complemented by high-resolution spatial transcriptomics using Visium HD, enabling the spatial mapping of all major skin compartments, including the epidermis, dermis, vasculature, immune and neuronal lineages, and appendages with emphasis on the PSU. We identify both well-characterized and previously underappreciated skin cell states, benchmark their spatial localization, and demonstrate robust integration through quantitative assessment using the scIB benchmarking framework. Consistent cell type nomenclature and metadata annotation further ensure downstream modeling, comparison, and interoperability.

**Fig. 1:**
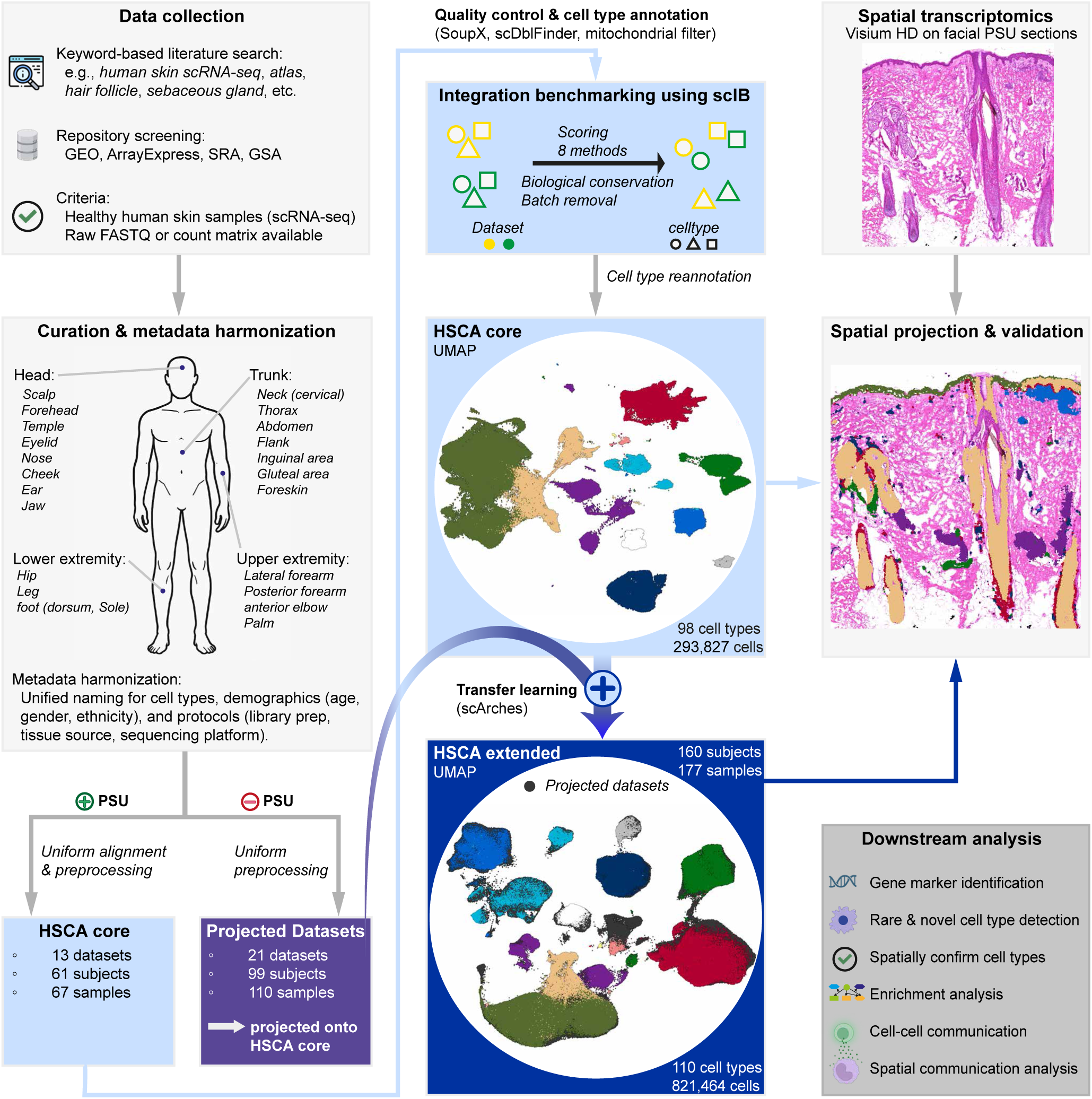
Study design integrating scRNA-seq and spatial transcriptomics for HSCA generation. Healthy human skin scRNA-seq datasets were collected and curated. Datasets were divided into PSU-containing and PSU-free samples. PSU-containing datasets underwent standardized reanalysis and processing, and integration performance was benchmarked. The most suitable tool was used to integrate these datasets into the HSCA core, followed by cell type annotation. Through transfer learning, 21 additional PSU-free datasets were incorporated, resulting in the HSCA extended (160 subjects, 177 samples, 110 cell types, >800,000 cells). Gene marker signatures were validated and refined using Visium HD spatial transcriptomics. Downstream analyses included the identification of novel and rare cell types, functional enrichment, and cell–cell communication analysis.

Our integrated skin atlas provides a comprehensive reference of the healthy human cutaneous landscape, enabling robust cell type annotation and offering a spatially resolved framework to investigate skin biology, disease-associated perturbations, benchmark organoids and in vitro models, and guide regenerative and precision medicine.

## 2 RESULTS

### Construction and integration of the HSCA core from PSU-containing datasets

We collected 34 publicly available scRNA-seq datasets of healthy human skin (Supplementary Table 1). A central objective of the HSCA is to resolve the cellular architecture of the PSU, which remains underrepresented in existing studies. To this end, we first constructed the HSCA core from datasets containing PSU cells.

Manual re-analysis of the datasets (see Methods) revealed that only 13 of the 34 datasets contained PSU cells, with often sparse representation (Fig. 2d). To ensure analytical clarity and biological relevance, we restricted the HSCA core to these 13 datasets^15,16,22–32^, providing the necessary resolution to define PSU-associated cell types while avoiding confounding complexity from datasets lacking PSU cells.

**Fig. 2:**
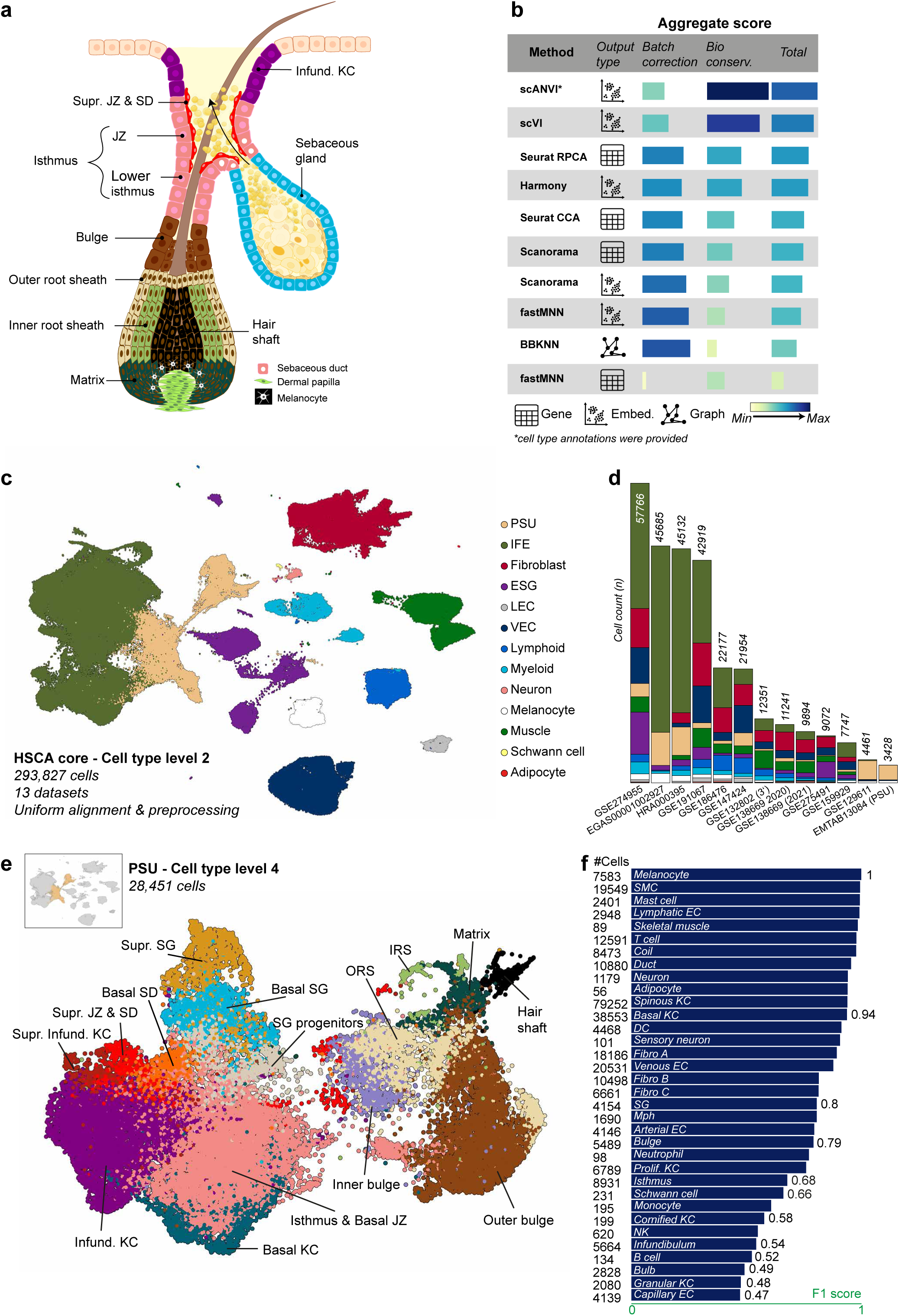
Construction and validation of the HSCA core. (**a**) Schematic illustration of the PSU. (**b**) Integration benchmarking using the scIB framework, showing aggregated scores for technical batch correction and biological conservation across methods. (**c**) UMAP representation of the HSCA core, integrating PSU-containing datasets with unified preprocessing and cell type annotation. (**d**) Composition of cell types across individual studies contributing to the HSCA core, as derived from the UMAP in c. (**e**), Zoom-in view highlighting the PSU cluster within the HSCA core. (**f**), Annotation performance of the HSCA compared to individual study-level annotations, quantified by F1 score. Abbreviations: see Supplementary Table 3.

To minimize technical variability, all raw sequencing reads in the HSCA core were reprocessed using a unified alignment and preprocessing pipeline (see Methods). Technical metadata (library preparation protocol, tissue sampling method, sequencing platform) and biological metadata (donor age, sex, ethnicity, anatomical site) were harmonized. Spatial information was captured in a three-tiered anatomical hierarchy (region, subregion, fine subregion), and cell type nomenclature was standardized to ensure consistent annotation (Supplementary Table 2), with abbreviations used for cell types listed separately (Supplementary Table 3).

To identify the optimal method for integrating the heterogeneous HSCA core datasets, we benchmarked multiple approaches using the scIB framework^21^ (Fig. 2b and Supplementary Fig. 1). All methods displayed a trade-off between batch effect removal and biological conservation, with scANVI^33^ and scVI^34^ performing best, consistent with previous reports^10,21^. However, biological conservation metrics rely on pre-existing cell type annotations within each dataset and therefore cannot capture novel or rare cell states that only emerge at the atlas level. Because scANVI uses these annotations as input and optimizes agreements with them, it may inadvertently remove genuine biological variation not represented in them^35^. In our context, where the goal is to discover novel or rare cell states, this dependence on prior annotations introduces a potential bias in its assessment using these metrics. By contrast, scVI operates fully unsupervised, while still providing strong correction of technical effects. Commonly used methods such as Harmony^36^ tended to overcorrect, obscuring biologically meaningful differences.

Applying scVI to the curated and reprocessed PSU-containing datasets yielded the HSCA core, comprising 293,827 cells (Fig. 2c). Cells were annotated across five hierarchical levels (Supplementary Fig. 2), with Level 1 representing broad lineages including cutaneous epithelial, stromal, immune, endothelial, and neural, and Level 5 capturing fine-grained states, such as Cortex cells of the PSU. At the finest resolution, the HSCA core atlas encompasses 98 distinct cell types. The PSU-focused subset, containing 28,451 cells, includes all major cell types of the PSU, providing comprehensive coverage of this key epithelial structure (Fig. 2e).

### Reference-atlas integration markedly raises cell type annotation fidelity

A key advantage of building a reference atlas is the improved accuracy of cell type annotations. To assess this quantitatively, we compared the dataset-specific annotations (manually assigned by us for each of the 13 datasets before integration) with the atlas-derived annotations that resulted from the HSCA core integration (Level 3 of our hierarchical taxonomy).

We constructed a confusion matrix across all cells (Supplementary Fig. 3) and calculated the F1-score to quantify the agreement between dataset-level annotations (treated as predictions) and the atlas annotations (treated as ground truth). The consolidated atlas annotations substantially outperform the initial labels for most cell types (Fig. 2f). High F1-scores (≥ 0.9) were observed for robust cell types, including melanocytes, smooth muscle cells, mast cells, and T cells. Notably, melanocytes achieved near-perfect agreement (F1 ∼ 1.0), highlighting the stability of this cell type across individual studies. By contrast, transitional or transcriptionally subtle populations exhibited lower F1-scores. These include granular (F1 = 0.48) and cornified keratinocytes (F1 = 0.58), as well as region-specific PSU cell types from the bulb (F1 = 0.49), isthmus (F1 = 0.68), and infundibulum (F1 = 0.54).

These low F1-scores highlight both the challenges of accurate cell type annotation in individual studies and the substantial improvement provided by the consolidated atlas, demonstrating its value as a reliable benchmark for future skin scRNA-seq datasets. Notably, annotation stability does not simply reflect cell numbers: populations such as melanocytes and skeletal muscle cells, despite moderate or low counts (e.g., 89 skeletal muscle cells), show highly segregating transcriptional features and robust annotation across datasets.

### Spatial validation of PSU architecture and hair-cycle dynamics with Visium HD

To validate the spatial organization of the PSU defined in our core atlas and to assess additional relevant cell types, we generated two 10X Visium HD spatial transcriptomics sections (8 µm spot diameter) derived from healthy facial skin of a 48-year-old White female donor (Fig. 3a, b). Both sections, designated D1 and D2, were obtained from the temporal region and each contained complete longitudinal PSUs. After standard Space Ranger processing, the maximum number of genes detected per spot was in 3,683 D1 and 3,199 in D2 (Fig. 3a and b). Spots were subsequently annotated based on marker gene expression, enabling robust identification of major skin cell types (Supplementary Fig. 4).

**Fig. 3:**
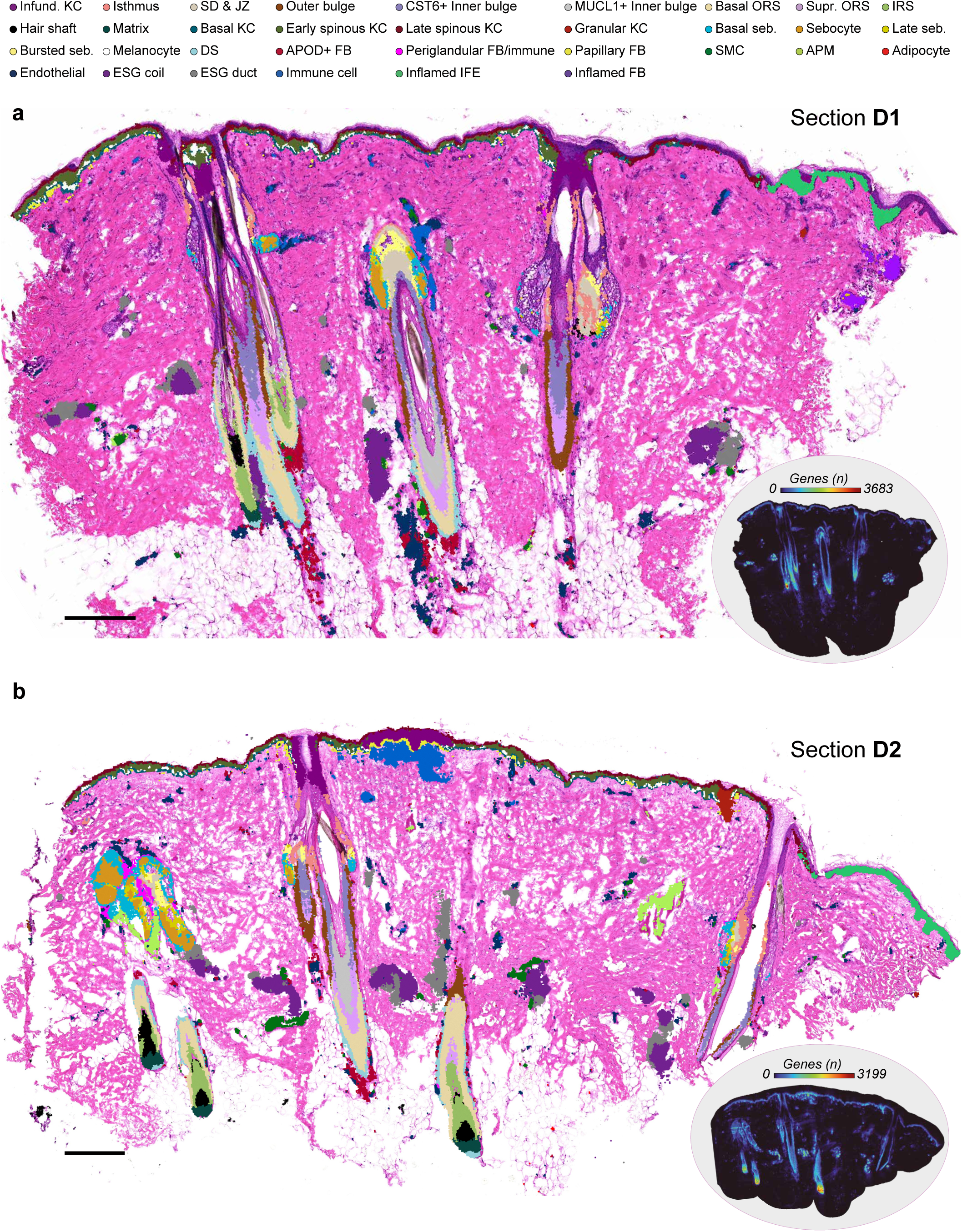
Spatial validation of PSU architecture with Visium HD. (**a**, **b**) Two 10X Genomics Visium HD spatial transcriptomic sections (8 µm spot diameter) derived from healthy facial skin of a 48-year-old White female donor (temporal region). Spots were annotated with marker gene expression, and the derived cell types are overlaid on the H&E sections. The bottom-right inset of each panel displays the number of detected genes per spot (maximum 3,683 in D1 and 3,199 in D2). Bar = 250 µm. Abbreviations: see Supplementary Table 3.

We performed a targeted subclustering of the Visium HD spots that overlapped the hair bulb. This enabled the characterization of major cell types within the bulb, including early and late matrix progenitors, basal outer root sheaths, companion layer, the three concentric inner root sheath layers (Henle’s, Huxley’s, cuticle), hair shaft cells (cortex and medulla), melanocytes, and dermal papilla fibroblasts (Fig. 4a, b).

**Fig. 4:**
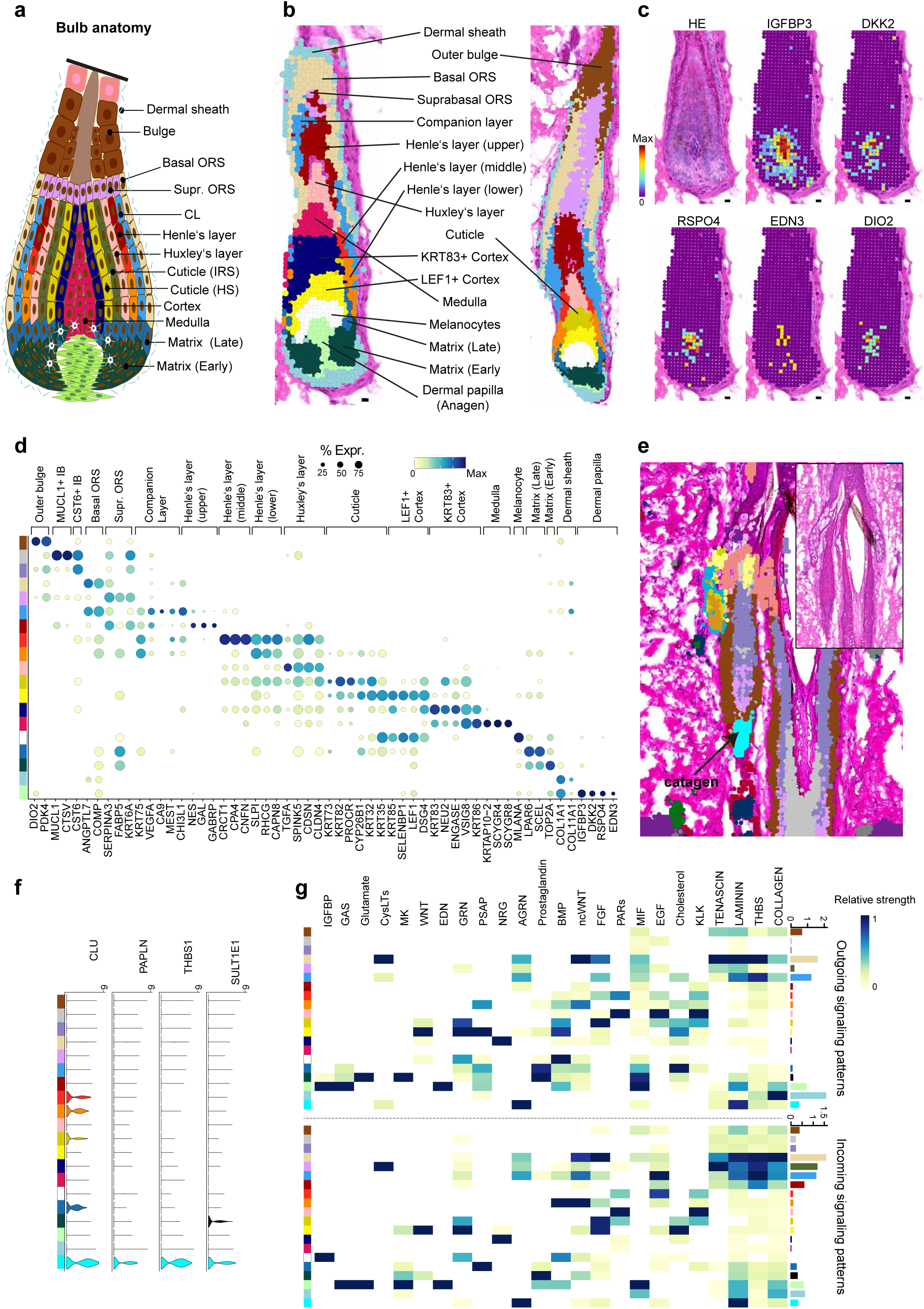
Spatial characterization and crosstalk of the human hair bulb using Visium HD. (a) Illustrative schematic of hair bulb anatomy. (b) Visium HD spots corresponding to the hair bulb overlaid on the tissue section. (**c**) Spatial feature plot of Dermal papilla markers. (**d**) Dot plot showing marker gene expression across major bulb cell types. (**e**) Catagen hair follicle section (D2) highlighting cell clustering. (**f**) Violin plots of gene expression in the catagen follicle cluster, reflecting hair-cycle-specific transcriptional dynamics. (**g**) Heatmap of spatial ligand-receptor crosstalk between follicular compartments inferred by CellChat. Bar = 8 µm. Abbreviations: see Supplementary Table 3.

Marker genes include *ANGPTL7* and *COMP* in the basal outer root sheath, *VEGFA* and *CA9* in the companion layer, *GAL* and *CRCT1* in Henle’s layer, *TGFA* in Huxley’s layer, *KRT73* in the cuticle, *LEF1* and *KRT83* in the cortex, keratin-associated proteins (*KRTAPs*) together with *SCYGR8* in the medulla, and *TOP2A* in the matrix cells. The Henle’s layer appears to be subdividable into upper, middle, and lower sub-regions, each suggesting a distinct transcriptional signature (e.g., *GAL* enriched in the upper zone, *CRCT1* in the middle zone, and elevated *RHCGG* in the lower zone) (Fig. 4d). Dermal papilla cells expressed specific markers including *IGFBP3*, *DKK2*, *RSPO4*, and *EDN3* (Fig. 4c and d).

Importantly, one of the analyzed follicles (section D2) was in catagen (Fig. 4e), enabling the investigation of hair-cycle specific transcriptional changes. In this regressing follicle, we observed expression of *SULT1E1*, *THBS1*, *PAPLN*, and *CLU*, reflecting the dynamic transcriptional landscape associated with hair-cycle progression (Fig. 4f).

### Spatial crosstalk analysis of follicular compartments with Visium HD

Unlike bulk or single-cell approaches, which either average out signals across compartments or lose anatomical context, our spatial crosstalk analysis using Visium HD enabled the assignment of ligand-receptor interactions to specific follicular compartments^37^. At this resolution, the composition of the human hair bulb has not previously been mapped, allowing us to chart intercellular communication within this structure for the first time (Fig. 4g). Dermal papilla fibroblasts emerged as prominent signal emitters, releasing *IGFBP*-, *GAS (Growth arrest-specific)*-, and *EDN*-mediated cues to neighboring cells. Matrix progenitors were the source of prostaglandin signals, whereas the medulla exhibited minimal communication, reflecting its terminally differentiated state.

Within the hair shaft, *KRT83*⁺ cortex cells preferentially engaged in neuregulin (*NRG*) signaling, while adjacent *LEF1^+^* cortex cells relied on canonical *WNT* cues, indicating a functional split between cortical subpopulations (Fig. 4g). The Huxley’s layer served as a source of *EGF*, whereas the lower Henle region acted as a strong receiver of *BMP* and *non-canonical WNT* (*ncWNT*) signals. Basal outer root sheath cells also displayed *ncWNT* activity. In the lower catagen bulge, the extracellular matrix protein *AGRN* dominated the outgoing network.

Collectively, these findings provide a spatially resolved framework for understanding how discrete follicular layers coordinate growth, regression, and terminal differentiation.

### Concordance and divergence of lower follicular compartments in HSCA and Visium HD

In the HSCA core, we identified 8,572 cells from the lower follicular compartments (Fig. 5a). Integrative comparison using RCTD decomposition^38^ revealed strong concordance of cell type gene signatures between the core atlas and Visium HD data (Fig. 5b). However, middle Henle’s layer and medulla cells were absent from the scRNA-seq atlas but clearly resolved in the Visium HD sections (Fig. 4b and d). Conversely, the HSCA contained a distinct cluster not captured in the Visium HD data, defined by a transcriptional profile consistent with the secondary hair germ (SHG) (Fig. 5c and d), a population previously described in murine telogen follicles^39^. The absence of SHG cells in the Visium HD dataset likely reflects the lack of telogen follicles in the analyzed sections (Fig. 3a and b).

**Fig. 5:**
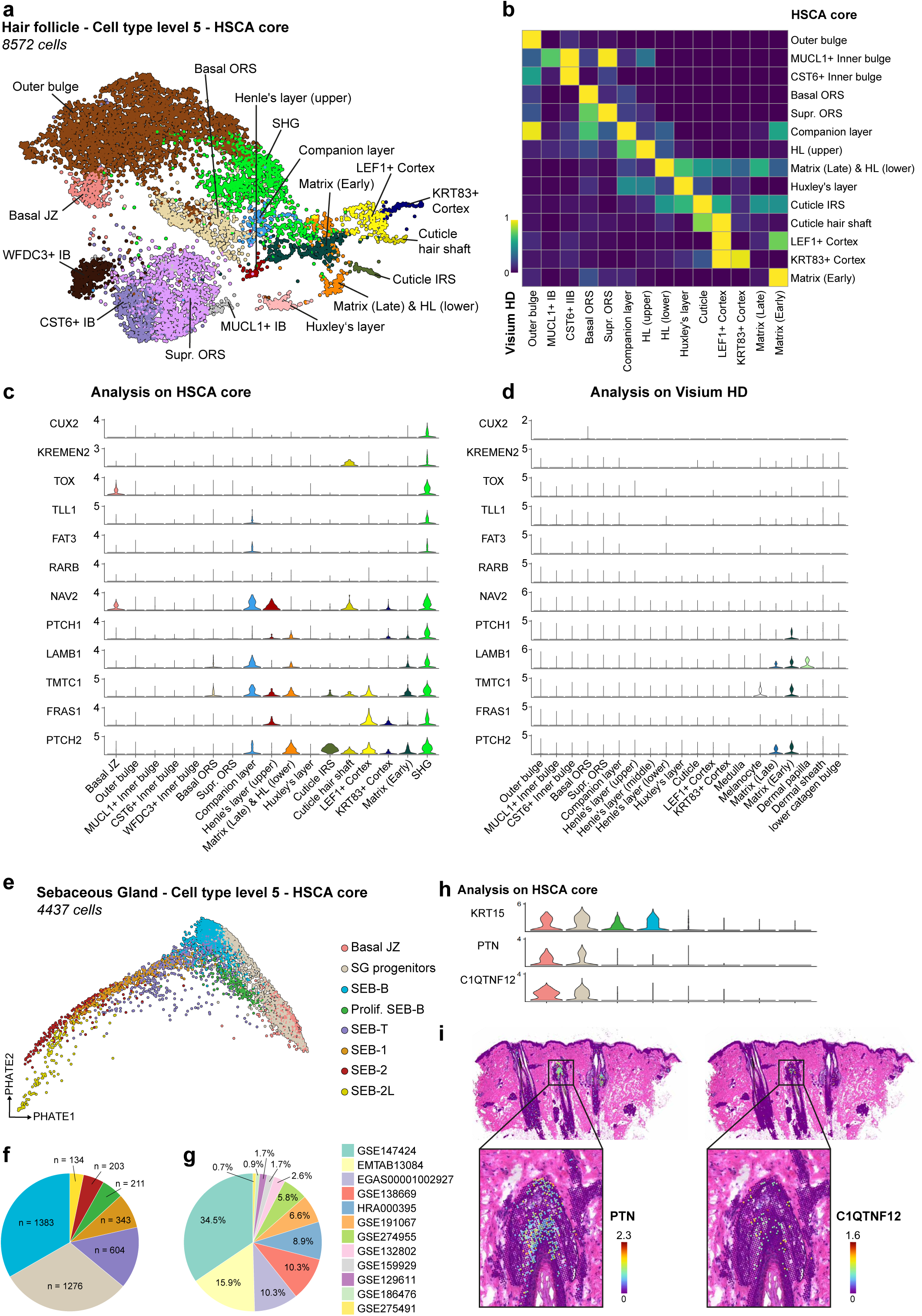
Concordance of lower follicular compartments in the HSCA core and Visium HD, and sebaceous gland differentiation trajectories. (**a**) UMAP of the HSCA core restricted to 8,572 cells from lower follicular compartments. (**b**) RCTD deconvolution of Visium HD data (from Fig. 4b) using the HSCA core, showing concordant cell type gene signatures. (**c**, **d**) Violin plots of marker gene expression for the SHG in the HSCA core (c) and in Visium HD (d). (**e**) PHATE embedding of sebaceous gland cells illustrating differentiation trajectories. (**f**) Pie chart summarizing the relative abundance of sebocyte maturation stages in the HSCA core. (**g**) Pie chart showing dataset origin of sebaceous cells across maturation stages. (**h**) Violin plots of *PTN* and *C1QTNF12* expression in sebaceous progenitors and the JZ in the HSCA core. (**i**) Independent spatial validation of *PTN* and *C1QTNF12* expression in Visium HD sections. Abbreviations: see Supplementary Table 3.

### Sebaceous gland differentiation trajectories in the HSCA core

We reanalyzed the sebaceous gland cluster from the HSCA core to resolve differentiation trajectories using PHATE^40^ (Fig. 5e). For comparison, a UMAP embedding is shown in Supplementary Fig. 5. The data recapitulate the sebocyte maturation stages described by Düz, et al. ^41^ (Supplementary Fig. 6). A key advance of our analysis is the resolution of the SEB-2L stage, which remained unresolved in the earlier scRNA-seq dataset^41^, underscoring the enhanced sensitivity of the HSCA core. In addition, the HSCA core corroborated that lipid metabolic programs initiate in SEB-1 and continue to be active in SEB-2, while also delineating additional distinct marker signatures that distinguish SEB-2 and SEB-2L from SEB-1 (Supplementary Fig. 6).

A pie chart summarizes the relative abundance of these maturation stages identified in the HSCA core (Fig. 5f). Notably, the proportion of cells representing late-stage sebocytes progressively declines along the trajectory. This underrepresentation likely stems from the biology of holocrine secretion^42,43^, as maturing sebocytes enlarge, accumulate lipids, and ultimately rupture, making them less likely to be captured by droplet-based scRNA-seq^44^. We also observed a pronounced enrichment of sebocytes in dataset GSE147424^25^ (Fig. 5g).

Moreover, sebaceous progenitor cells and the junctional zone display strong enrichment for PTN and C1QTNF12 (Fig. 5h), both of which are spatially validated in Visium HD sections, highlighting their role as early lineage markers (Fig. 5i) and showing how high-resolution spatial transcriptomics enables precise mapping of cell states^45^.

### Discovery of a CCER2⁺ Merkel cell cluster

By integrating the HSCA core, we identified a previously unrecognized cluster that was not detectable in any individual dataset (Fig. 6a). This cluster is characterized by strong expression of CCER2 (Fig. 6b), a gene that is poorly characterized overall^46^ and, to our knowledge, has not been reported in skin. In addition, the cluster shows co-expression of KRT20 (Fig. 6b), a well-established marker for Merkel cells^47,48^. VIP was also detected in Merkel cells^49^.

**Fig. 6:**
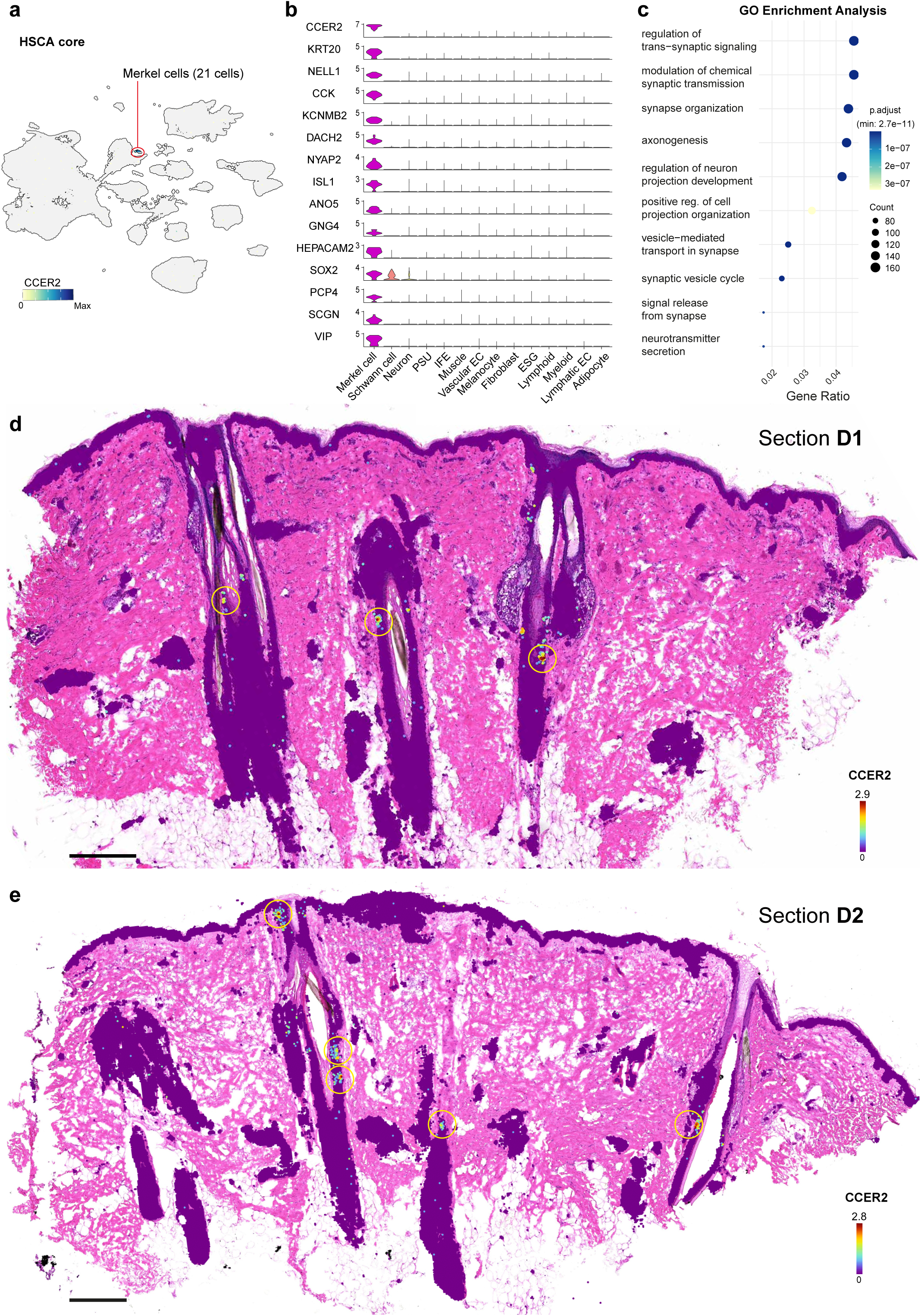
Identification and characterization of Merkel cells using the HSCA core. (a) Feature plot of *CCER2* expression highlighting the Merkel cell cluster in the HSCA core. (b) Gene signature of the cluster, including the characteristic *KRT20* marker for Merkel cells. (c) Functional enrichment analysis of the Merkel cell gene signature, visualized as a dot plot. (**d**, **e**) Spatial visualization of *CCER2* expression in the bulge region of hair follicles in Visium HD sections.

Merkel cells are specialized neuroendocrine cells involved in tactile perception, whose abundance varies across body sites but is highest in the epidermis of the lips and fingertips^50^ as well as within hair follicles^51–53^. Using Visium HD, we localized these cells mainly to the hair follicle bulge region in our dataset, a known stem cell reservoir (Fig. 6d and e). In the D2 section representing catagen-stage follicles, they were additionally detected in the infundibulum (Fig. 6e). The gene signature shown in Fig. 6b was further visualized as a module score in Supplementary Fig. 7.

The presence of this CCER2⁺ Merkel-cell population, overlooked in previous skin studies, underscores the enhanced sensitivity of the HSCA core and suggests a potential link between mechanosensation and the bulge stem cell niche.

### Construction of the extended Human Skin Cell Atlas via transfer learning

The HSCA core was built from studies that contain PSU cells and provide raw files so that we could apply a uniform preprocessing pipeline. A total of 21 scRNA-seq datasets^14,22,24,27,54–69^ did not meet these criteria: 20 lacked PSU cells, and one dataset (GSE179633^54^) included PSU cells but did not provide raw data. A systematic metadata audit of all 34 datasets (Fig. 7a) revealed that essential donor information (sex, age, ethnicity and anatomical site) is missing for a substantial fraction of samples, whereas technical metadata (platform and sampling method) is almost always present^11^. This paucity of biological metadata limits population-level analyses such as age- or ethnicity-specific comparisons. To nevertheless leverage the biological signal contained in these datasets, we employed scArches^35^, a transfer learning framework that maps new datasets onto a reference without retraining the entire model (see Methods). This enabled us to project these datasets onto the HSCA core and construct the extended Human Skin Cell Atlas (HSCA extended), comprising 160 donors, 177 samples, 110 distinct cell types, and 821,464 cells (Fig. 7b). In the resulting UMAP, HSCA core cells are highlighted within the broader landscape of the extended dataset, which is shown in black (Fig. 7b). scArches also enables cell type label transfer from the HSCA core to the HSCA extended using a k-nearest neighbor graph and provides an uncertainty score for each assignment^35^. Label transfer uncertainty is shown in blue on the UMAP in Figure 7c. As described previously^10^, uncertainty naturally accumulates in transitional zones between related states. Regions of very high uncertainty, which overlap with the black areas in Figure 7b, correspond to novel cell populations absent from the HSCA core, including chondrocytes, neural progenitors, erythrocytes, and ex vivo cultivated Schwann cells (Fig. 7d and Supplementary Fig. 8). Even within fibroblasts and PSU cells, which are represented in the HSCA core, elevated uncertainty is observed due to the presence of closely related functional states.

**Fig. 7:**
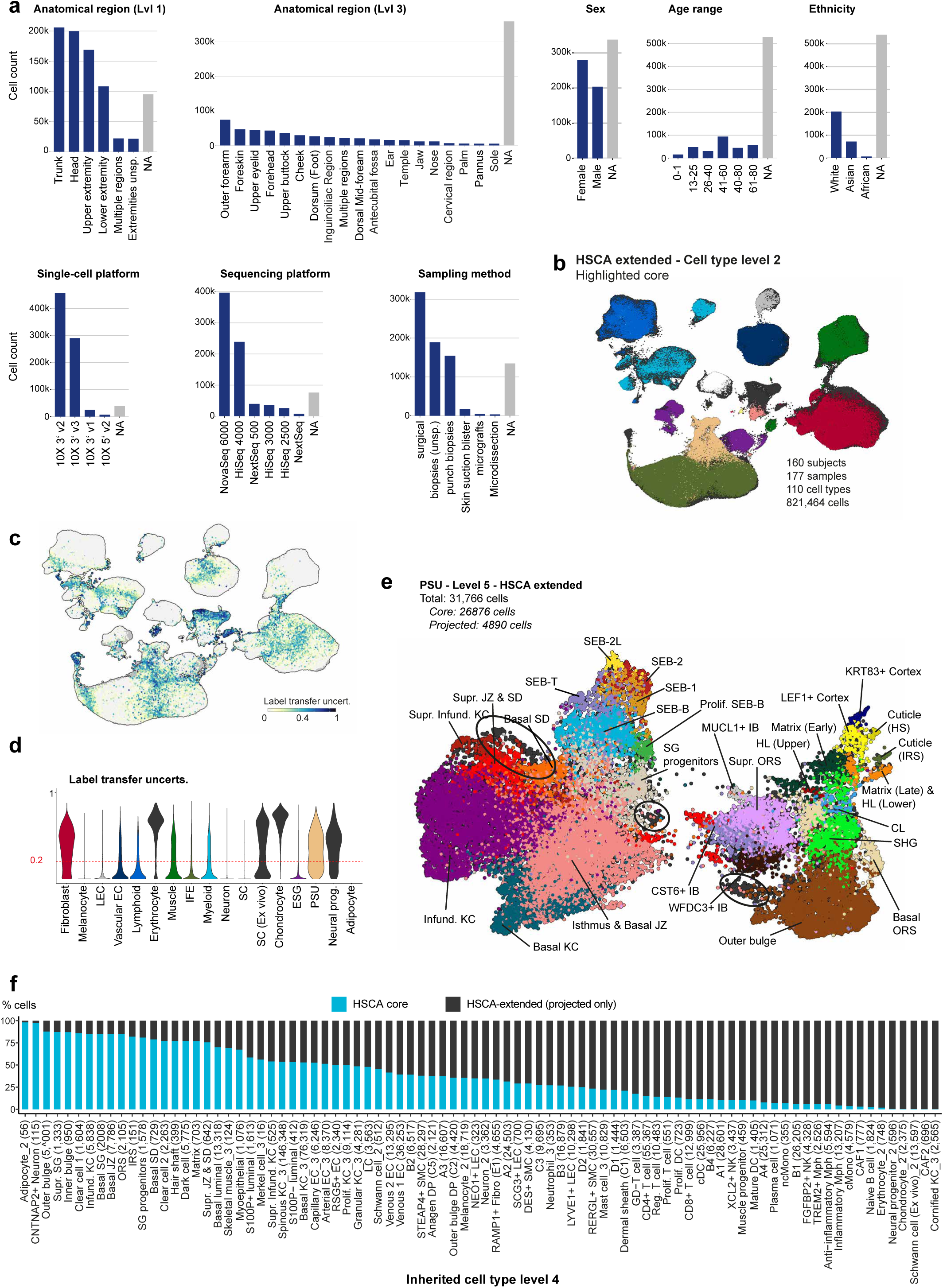
Construction and characterization of the HSCA extended via transfer learning. **(a)** Histograms summarizing the availability of key metadata fields across samples. (**b**) UMAP of the HSCA extended (821,464 cells) with HSCA core cells highlighted in color and projected datasets shown in black. (**c**) Label transfer uncertainty of HSCA core cell type annotations projected onto the extended dataset, visualized on the UMAP. (**d**) Label transfer uncertainty grouped by cell type (level 2) and aggregated from level 5 annotations, highlighting areas of elevated uncertainty. (**e**) UMAP of the PSU extended, with HSCA core cells highlighted, showing that no relevant variance is introduced for PSU lineages. The circles highlight extension-specific cell variations that are not found in the HSCA core. (**f**) Proportional contribution of each cell type, showing the fraction derived from HSCA core versus HSCA extended at inherited cell type level 4. Abbreviations: see Supplementary Table 3.

Because label transfer uncertainty highlighted gaps in the HSCA core, we performed a de-novo re-annotation of the entire HSCA extended (Fig. 7e and Supplementary Fig. 8-15). Overlaying the PSU core on the projected datasets shows that no additional biological variance is gained for PSU lineages (Fig. 7e and Supplementary Fig. 9), which was expected since the HSCA core was deliberately built by selecting PSU-containing cells. In contrast, other lineages benefit substantially: the extended dataset not only increases cell numbers for existing populations but also reveals new cell types (Fig. 7f). Supplementary Figure 10 shows the HSCA core highlighted in color on the extended data for immune cells, illustrating strong concordance in cluster assignments. Not only does the number of data points per cluster increase, but novel cell types absent from the HSCA core are also identified, including naive CD8^+^ and CD4^+^ T cells, a finer subdivision of naive B cells and plasmacytoid dendritic cells, as well as greater variance within classical monocytes (Supplementary Fig. 10). Fibroblast coverage also expands (Supplementary Fig. 11). Leveraging the marker sets reported by Ascension, et al. ^70^ and Ascensión and Izeta ^20^, we observe clearer separation of fibroblast submodules along axis A, which were previously blurred in the HSCA core due to limited cell numbers (Supplementary Fig. 11). Among the 177 samples, a single sample (GSM5050540^57^) unexpectedly harbors two distinct cancer-associated fibroblast (CAF) clusters (RGS5⁺, MCAM⁺, NOTCH3^+^, CHI3L1⁺, TMEM158⁺)^71,72^ that together represent ∼0.8 % of the cells (∼1473 cells) (Supplementary Fig. 16). This observation underscores that rare disease-associated populations can be present even in samples labeled as healthy. In addition, we analyzed IFE, eccrine sweat-gland, muscle, and endothelial populations, and assessed their concordance with the HSCA core (Supplementary Fig. 12-15).

In summary, scArches-based transfer learning enables rapid, first-draft integration of heterogeneous skin datasets, provides per-cell uncertainty scores for label transfer that guide downstream de novo re-annotation, and demonstrates both the sufficiency of the HSCA core for PSU-lineage representation and the additional biological variance gained when the atlas is extended to the full public repertoire.

## 3 DISCUSSION

We present the human skin cell atlas (HSCA), integrating 34 scRNA-seq datasets from 160 donors and complemented by high-resolution spatial transcriptomics. The HSCA resolves major skin compartments and appendages, including the PSU, with hierarchical annotations from broad cell types to finely resolved subpopulations. While the HSCA core captures most skin cell types, the extended dataset enhances statistical power for rare populations and refines transitional states between closely related lineages.

The PSU has been underrepresented in previous omics studies due to its rarity, uneven spatial distribution, and difficulty in capturing keratinized follicular cells and large sebocytes. Targeted microdissection increased PSU cell numbers (Fig. 2d), but scRNA-seq loses tissue context. To overcome this, we complemented scRNA-seq with Visium HD, resolving PSU architecture at unprecedented granularity. However, Visium HD only captures structures present in the section, so features such as the putative secondary hair germ identified by scRNA-seq integration were not detected (Fig. 5c and d). Furthermore, the curved follicle architecture often caused loss of upper PSU cells due to difficulty capturing all cells in one plane (Fig. 3a and b). While permeabilization was consistently effective in lower follicle compartments, sebaceous glands showed variable results, even within the same section, likely reflecting structural or biochemical differences, which could be addressed by compartment-specific protocols^41^.

Our data recapitulate known signaling axes in hair biology, particularly those centered on DP cells^73–77^. Prostaglandin signaling, a key regulator of hair growth^78,79^, was spatially assigned to matrix cells. Canonical WNT signaling, which governs hair fate decisions^80–82^, localized to LEF1^+^ Cortex cells activity, consistent with its role in anagen entry^83^. Beyond these established pathways, we anatomically mapped underexplored signals in hair biology. NRG signaling, implicated in epidermal and follicular development^84^, localized to cortex cells, whereas EGF/TGF-α signaling, was here strongly secreted by Huxley’s layer. Finally, we identified Agrin-mediated signaling in the bulge during catagen, suggesting a role in epithelial-mesenchymal remodeling and regenerative niche turnover^85^.

A key advance of the HSCA is that it enables systematic evaluation of how library preparation protocols affect cell type recovery. For example, one dataset^25^ shows a marked enrichment of sebocytes in extremity skin, exceeding even PSU-rich scalp datasets. This enrichment was confined to frozen samples, suggesting that freezing may better preserve this fragile cell type. Such protocol effects could be critical for acne research, where scRNA seq datasets that robustly capture sebocytes are lacking^86,87^. Conversely, a 48-h ex vivo explant protocol without enzymatic digestion enabled recovery of stratum corneum keratinocytes^64^ (Supplementary Fig. 12), usually lost in standard workflows, highlighting a simple way to capture fragile, anucleate corneocytes. Because the HSCA is anchored in native tissue, it also provides a benchmark for evaluating in-vitro skin models and organoids. As a proof-of-concept, cultured Schwann cells mapped only weakly to their in-vivo counterparts. Similarly, the abundant sebocytes offer a reference for comparing immortalized lines such as SZ95^88^.

In the HSCA core we uncovered a rare KRT20⁺/CCER2⁺ Merkel-cell population that had not been reported in any individual dataset or in previous studies. Our spatial mapping placed these cells in the hair-follicle bulge, a stem-cell reservoir^89^, and transiently in the infundibulum of a catagen-stage follicle, hinting at a hair-cycle-dependent migration. Notably, adding 21 PSU-absent datasets did not enlarge the cluster, reinforcing the bulge as the dominant niche. Understanding how Merkel cells interact with bulge stem cells could open new avenues for hair-follicle regeneration. If CCER2-mediated signalling modulates stem-cell activation or quiescence, pharmacological manipulation of this pathway might be exploited to re-activate dormant follicles in alopecia.

A consistent, community-endorsed cell-type nomenclature is essential. We therefore adopted a provisional naming scheme based on literature and our annotations, intending the HSCA to evolve through expert consensus. Metadata harmonization was a major effort, yet several public datasets lacked complete sample information, highlighting the need for community-wide metadata standards. Future HSCA releases will expand metadata coverage and enabling more robust analyses of biological covariates.

Looking ahead, the HSCA provides an ideal resource for the development of skin-specific foundation models^9^. Inspired by advances in AI, such models learn generalizable representations from diverse datasets and enable various downstream tasks with minimal fine-tuning. For skin research, foundation models could accelerate cross-dataset cell type annotation, gene regulatory network inference, and perturbation analysis, shifting the focus from descriptive atlases to functional models^90^.

Achieving this requires integrating additional modalities, such as scATAC-seq to map regulatory elements^91^, CITE-seq to profile functional protein effectors^92^, and snRNA-seq to better capture fragile or dying cells^93^. Furthermore, including diseased skin is essential to benchmark altered skin biology, discover novel cell populations and markers, and resolve site-specific disease patterns in conditions like acne and psoriasis^10,11,94^. However, integrating these diverse datasets poses challenges^9,21^, such as the need for comprehensive reannotation and the mitigation of batch effects.

Together, these efforts will ensure the HSCA evolves into a comprehensive, multi- modal resource, enabling predictive and functional models of skin biology in health and disease^95^.

## ETHICS STATEMENT

The Visium HD study adhered to the ethical standards of the Declaration of Helsinki. Facial skin explants were obtained from a healthy donor undergoing aesthetic surgery, with Alphenyx providing the tissue. All necessary approvals were obtained by Alphenyx from the French Ministry of Research and approved by the French Ethical Committee (n° AC-2019-3567 and IE-2019-1077). Documentation confirming ethical approval is available upon request. Written informed consent was obtained from all adult participants prior to tissue collection.

## MATERIALS AND METHODS

### Experimental design of wet laboratory studies

#### Subject details

Full-thickness human facial skin specimens from the temporal region were obtained from a healthy, 48-year-old White female undergoing facelift surgery, with written informed consent. The donor had a BMI of 22.2, low sun exposure, was a non-smoker, non-menopausal, used no hormonal contraception, and had no signs of baldness. Subcutaneous fat was removed, and the skin was cut into 6 × 8 mm pieces, embedded in OCT mounting medium, and frozen at –80 °C for seven months prior to sectioning for Visium HD analysis.

#### Visium HD Spatial Transcriptomics

Fresh frozen (FF) human skin tissue was processed for Visium HD spatial transcriptomics following 10x Genomics guidelines. All experimental steps were performed by Crown Bioscience Germany GmbH.

### Tissue Sectioning and RNA quality control

The FF skin tissue block was sectioned using a cryostat (Leica): Three 5 µm sections were used as curls for RNA isolation and quality control and one 5 µm section was mounted on Schott Nexterion™ 3-D Hydrogel (H) coated slides (Fisher Scientific) for the Visium HD Spatial Gene Expression workflow. RNA was isolated using the RNeasy

Plus Mini Kit (Qiagen) and quality assessed with an Agilent Fragment Analyzer, using the High Sensitivity RNA Kit (Agilent). Samples with RNA quality number (RQN) ≥ 4 were used for further analysis.

### Tissue preparation and 10x Genomics Visium HD Spatial Gene Expression Workflow

Tissue sections were formaldehyde fixed, H&E stained, scanned with Axio Scan.Z1 (Zeiss) and destained followed by tissue permeabilization according to manufacturer’s instruction (Visium HD FF Tissue Preparation, CG000763).

Library preparation was performed with the 10x Genomics Visium HD Spatial Gene Expression Reagent Kits and CytAssist to the user guide (10x Genomics, Visium HD Spatial Gene Expression Reagent Kits User Guide, CG000685). After permeabilization, an overnight whole-transcriptome hybridization with the human probe set v2.0 (18,536 protein-coding genes, 54,018 probes) was performed, followed by probe ligation and washes. Ligated probes were transferred to the Visium HD slide using CytAssist (v2.2.0.8), where release from tissue, capture, and extension to generate spatially barcoded ligation products occurred. These products were then pre-amplified, purified, and used for library construction.

### Sequencing and Alignment

Library was quality controlled and sequenced at a concentration of 2 nM on a NovaSeq® X Plus using a 10B, 300 cycle flowcell (Illumina).

Raw sequencing data were processed with Space Ranger (v3.1.3, 10x Genomics) for demultiplexing, alignment to the GRCh38-2024-A reference, trimming of reads, de-duplication, filtering, and generation of feature-barcode matrices, as well as aligned to microscopic images with the Image Reorientation set “on” and Filter probes set “on” resulting in 18,085 human protein coding genes targeted by 53,504 probes as filtered output for further spatial analysis.

### HSCA core raw data processing

#### Identification and acquisition of public human-skin scRNA-seq datasets

We systematically collected publicly available scRNA-seq datasets of healthy human skin through keyword searches in PubMed and Google Scholar. Peer-reviewed articles and preprints were screened for links to primary repositories (GEO, ArrayExpress, SRA/ENA, GSA, Human Cell Atlas), and only studies describing unperturbed healthy skin were included. For studies containing both healthy and diseased samples, only the healthy samples identifiable from metadata were retained.

For each accession, we examined whether a gene-by-cell count matrix was available. If such a matrix was deposited, it was downloaded directly. When only raw FASTQ files were available, reads were retrieved and re-processed using a uniform pipeline (see “Development of a core human skin atlas enriched for PSU cells”). We collected 30 publicly available scRNA-seq studies of healthy human skin (Supplementary Table 1). As individual studies often included multiple experimental conditions or sampling strategies, we subdivided them into distinct datasets:

- **Kim, et al.** ^27^ was split split into three-prime and five-prime datasets.
- **Ganier, et al.** ^24^ generated a full-thickness skin biopsy dataset (Ganier_Lynch_2024_1) and a separate microdissected PSU dataset from healthy scalp (Ganier_Lynch_2024_2).
- **Liu, et al.** ^14^ comprised trunk (Liu_Landen_2025_1; accession GSE241132) and foot (Liu_Landen_2025_2; accession GSE265972).
- **Cheng, et al.** ^22^ included samples from foreskin, breast, abdomen, and scalp.

Because foreskin lacks PSU structures, these samples were excluded from the HSCA core, designated as a separate dataset (Cheng_Cho_2018_2) in the Extension, and subsequently projected onto the HSCA core via transfer learning.

This workflow yielded 34 publicly available scRNA-seq datasets from healthy human skin, comprising 821,464 cells after quality control (Fig. 1).

#### Curation and harmonization of study-level metadata

Study-specific information, including donor characteristics, tissue provenance, sequencing chemistry, and technical parameters, was extracted from publications, supplementary information, and repository meta-tables. For each of the 34 scRNA-seq datasets, relevant fields were compiled into a master spreadsheet (Supplementary Table 1). Missing values were encoded as “NA” in the final atlas. Terminology was harmonized across studies to produce a unified, machine-readable metadata table that supports cross-study comparability and downstream modeling. A complete description of all metadata fields is provided in Supplementary Table 4.

#### Development of a core human skin atlas enriched for PSU cells

To create a high-quality PSU core atlas, all identified scRNA-seq datasets were screened for PSU cell types based on expression of well-established PSU marker genes, using either the deposited count matrices or count matrices generated from raw sequencing data.

Datasets containing PSU cells (n=13) were reprocessed from raw data. In most cases, this consisted of FASTQ files deposited in public repositories. For two accessions (GSE138669^28^ and EGAS00001002927^22^), only BAM files were available, from which FASTQ files were generated using the bamtofastq functionality from Cell Ranger^96^.

All raw data were uniformly aligned to GRCh38 using Cell Ranger v7.2.0 (default parameters) to generate gene-by-cell count matrices, ensuring comparability and avoiding alignment-related bias.

#### Processing of count matrices

Filtered feature-barcode matrices generated from the aligned raw data (see “Development of a core human skin atlas enriched for PSU cells”) were imported into R, and individual Seurat objects were created with annotated sample metadata. Ambient RNA contamination was estimated and corrected using SoupX^97^. For each sample, clusters were first identified after normalization, variable feature selection, scaling, and PCA-based dimensionality reduction. These cluster assignments were used as input for SoupX to model and adjust for ambient RNA contamination. Contamination fractions were automatically estimated unless visual inspection of the ambient-gene profile indicated otherwise (for datasets GSE191067^32^ and GSE274955^16^, a fixed contamination fraction of 0.1 was applied).

Doublets were identified using scDblFinder^98^, with singlets retained for all subsequent analyses. A lenient initial filtering step removed only cells with extremely low coverage (<200 reads) to avoid errors during doublet detection. Doublet detection was then performed on raw counts prior to stringent QC filtering, preventing underestimation of the expected doublet rate and enabling detection of doublets formed between high- and low-quality cells. After doublet removal, cells with more than 25% mitochondrial gene content or fewer than 200 detected genes were excluded.

Data were normalized, highly variable genes identified, scaled, and subjected to PCA. Batch effects were corrected using Harmony with sample identity as covariate^36^. The harmonized embeddings were used for UMAP visualization, neighborhood graph construction, and clustering. Subclustering within major cell categories refined annotations, and low-quality or ambiguous clusters were excluded from the final atlas.

#### Benchmarking pipeline for atlas integration

We applied the benchmarking pipeline from Luecken, et al. ^21^ (adapted to the objectives of the skin atlas). As input, we used a merged, preprocessed AnnData object comprising 13 HSCA core datasets. The pipeline performed feature selection (top 2000 highly variable genes) and scaling.

We compared integration methods including BBKNN, FastMNN, Harmony, Scanorama, scVI, scanVI, and Seurat (CCA and RPCA), which produced either integrated graphs, joint embeddings, or corrected feature spaces.

Integration performance was evaluated using the batch correction and biological conservation metrics described in Luecken, et al. ^21^. Batch correction metrics comprised ASW (batch), graph iLISI, kBET, graph connectivity, and principal component regression. Biological conservation metrics (label-dependent) included ASW (cell type), NMI, ARI, graph cLISI, isolated label F1, and isolated label silhouette. Metric values were averaged into batch and biological category scores, and combined into an overall score (weighted 40% batch, 60% biological).

The workflow was implemented in Snakemake, with each method executed in isolated environments to avoid dependency conflicts. Outputs included integrated UMAPs, metric score plots, and benchmarking plots to assess scalability.

#### Subclustering and bottom-up strategy for cell type annotation

The HSCA adopts a bottom-up hierarchical annotation workflow that differs from the typical top-down pipelines used in previous atlases^10^. In a top-down approach the integrated dataset is first projected into a low-dimensional space (commonly a UMAP of all cells) and clustered globally. Because the first clustering step is driven by the most abundant or highly variable transcriptional programs, dominant lineages can mask the signal of rare or transcriptionally subtle populations, leading to blurred cluster boundaries and occasional misassignments (e.g., non-epithelial immune cells grouped with interfollicular keratinocytes).

This problem is avoided in the HSCA’s bottom-up approach, which begins analysis at the finest level within broader cell type categories (often referred to as subclustering), resulting in clear cluster boundaries among related cell types. Once the fine annotation of a cell is correct, its identity can be unambiguously propagated to higher levels, which guarantees that the same cell cannot be assigned to contradictory lineages at different levels (Supplementary Fig. 2a). In some cases, such as mast cells, the most granular identity is reached at an intermediate level (e.g., Level 3). Here, the label is inherited to lower levels and annotated with a numeric suffix to indicate the level at which the classification was defined (Supplementary Fig. 2b).

Cell type annotation was based on marker genes identified in our dataset, which were anchored and validated using canonical and previously reported markers (see^10,14,18,20,30,41,75,99–103^).

#### Intuitive visualization of sebocyte differentiation with PHATE

PHATE visualization of the sebaceous gland was performed in R using the Python implementation accessed via the reticulate package. PHATE has been demonstrated to efficiently capture differentiation trajectories^40^. Default PHATE parameters were used except for the nearest-neighbor (knn) and principal-component (npca) settings, which were fixed at 30 to ensure that the PHATE embedding was directly comparable to the UMAP embeddings generated with Seurat’s default settings (Supplementary Fig. 5).

#### Merkel cell enrichment analysis

Differentially expressed genes in the Merkel cell cluster (HSCA core) were identified using Seurat’s FindMarkers function (Fig. 6b). Genes with a p-value < 0.05 were considered significantly differentially expressed and subjected to Gene Ontology enrichment analysis using the enrichGO function^104^. The analysis employed the human reference database org.Hs.eg.db, with gene symbols as identifiers, and focused specifically on biological processes. The top ten enriched terms were visualized (Fig. 6c).

### HSCA extended

#### Processing of HSCA extended query datasets

We added 21 publicly available scRNA-seq datasets of healthy human skin to the HSCA collection (Supplementary Table 1). Preprocessing followed the same pipeline and QC thresholds as applied for the HSCA core (see “Processing of count matrices”), with two exceptions. First, uniform re-alignment was not performed for all datasets because raw sequencing files were unavailable for several studies due to privacy restrictions. Instead, we used the feature-barcode matrices provided by the authors. Raw file availability and re-alignment status are listed in Supplementary Table 1. Second, ambient RNA correction was omitted, as this procedure requires raw gene-by-cell matrices directly obtained after alignment.

Processed Seurat objects were saved without embeddings (PCA, neighborhood graph, UMAP), as they serve solely as query data for transfer learning onto the HSCA core atlas.

#### Transfer Learning with scArches

Transfer learning (TL) leverages a model trained on a large, well-characterized single-cell reference to accelerate analysis of new query datasets^35^. In single-cell atlases, TL updates only a subset of parameters (adapter weights) while keeping the HSCA core network frozen. This approach reduces the amount of data required for integration, the computational time, and enables mapping of new datasets onto an existing reference^35^.

We first built a reference model using the HSCA core (Fig. 1). The reference was instantiated as a scVI model following the official scArches scVI vignette (<u>Unsupervised surgery pipeline with SCVI - scArches documentation</u>): three hidden layers (instead of two), covariate encoding enabled, and layer normalization applied to both encoder and decoder (batch normalization disabled). The model was trained for a maximum of 200 epochs using default parameters, with weight decay set to zero and early stopping enabled. The resulting scVI object was saved for mapping of query datasets.

To map 21 external healthy skin datasets, we applied the scArches *surgery* workflow, adapting the HLCA extended atlas pipeline. Each query matrix was subset to the 2,000 HVGs used in the reference. A sample-specific batch label (adapter) was inserted into the frozen network, and only adapter weights were updated. Adapter training was performed for a maximum of 200 epochs. Early stopping was triggered when the validation unweighted loss did not improve for 20 consecutive epochs, and the learning rate was reduced by a factor of 0.1 after 13 epochs without improvement.

Following surgery, the reference embedding and AnnData object were saved, converted to RDS, and used for de novo integration of selected cell populations, with uncertainty scores guiding bottom-up annotation (see “Cell type label transfer from HSCA core to extended datasets” and “Subclustering and bottom-up strategy for cell type annotation”). During this inspection, two samples (GSM8238437 and GSM8238438) displayed a wound-healing transcriptional signature, despite being originally labeled as healthy. These samples were therefore omitted from the final HSCA extended annotation.

#### Cell type label transfer from HSCA core to extended datasets

Cell type annotations from the HSCA core were propagated to HSCA extended datasets using the k nearest neighbour (kNN) routine in scArches (see “Mapping data to the Human Lung Cell Atlas” vignette). The latent coordinates of reference and query cells were concatenated to build a joint embedding, from which a kNN graph (*k* = 50) was constructed. Each query cell was assigned the most prevalent reference label among its neighbours, and an uncertainty score was computed^10,35^. Cells with an uncertainty score above 0.20 were labelled unknown, often representing rare, transitional, or populations absent in the HSCA core^10^. In total, the uncertainty scores guided de novo, bottom-up integration, enabling refinement of HSCA extended annotations (Supplementary Fig. 8-15).

### Visium HD downstream analysis

#### Whole-Skin analysis with Visium HD

Spatial transcriptomics data from two Visium HD skin sections (D1 and D2) were processed with Seurat^105^ and BANKSY^106^. Raw 10x Genomics outputs were imported using Load10X_Spatial, and the 8 µm assay was set as default. Spots expressing more than 100 genes and fewer than 25 % mitochondrial reads were retained. After QC, the two objects were merged and normalized via SCTransform^107^ to generate variance-stabilized expression values, used for downstream analyses.

Centroid coordinates were extracted and appended to the metadata for BANKSY. Dimensionality reduction with BANKSY was performed on the *SCT* data using lambda = 0.2 (spatial regularization), split.scale = TRUE (independent scaling per section), k_geom = 15 (geometric neighborhood size), and *group* = “Sample” to retain sample identity. The BANKSY latent space was stored in a new assay.

Principal component analysis was performed on the BANKSY assay (RunPCA, 30 PCs), and the PC embeddings were batch-integrated across the two sections using Harmony (RunHarmony, grouping variable = Sample). The Harmony-corrected dimensions were then used to compute a UMAP embedding (RunUMAP, dims = 1:30) and a shared nearest-neighbor graph (FindNeighbors). Clustering was carried out with FindClusters on the BANKSY reduction, yielding fine-grained spatial domains. Gene expression patterns were visualized directly on tissue sections (SpatialFeaturePlot) or embeddings (FeaturePlot), and cell type identities were assigned accordingly. Final visualizations were generated with TissUUmaps3^108^.

#### Processing lower hair follicle cells with Visium HD

Lower hair follicle cells were processed analogously to “Whole-Skin analysis with Visium HD”, but restricted to specific regions to achieve higher-resolution mapping (Fig. 4b). BANKSY dimensionality reduction used the same parameters, except for lambda = 0.15 and k_geom = 5; all other settings remained identical.

#### Cell–cell communication analysis with spatial context using CellChat

Cell–cell communication was analyzed with CellChat v2 following the standard recommended workflow, with default parameters unless otherwise specified^37^. For each spatial dataset, a CellChat object was created from the normalized expression matrix using createCellChat. Pixel-to-micrometer conversion was specified via spatial.factors: for D1, we set ratio = 0.644 (spot.size/spot_diameter_fullres), and for D2, ratio = 0.766. The tolerance parameter tol was set to spot.size/2 = 4 to account for variability when calculating cell-to-cell distances^37^.

We used the extended CellChat v2 database but excluded all cell–cell contact interactions, as these were highly overrepresented in our skin samples and could obscure the analysis of paracrine signaling pathways. Communication probabilities were computed with computeCommunProb (trim = 0.10). The contact.range was set to 25 µm, while the broader interaction.range for paracrine signaling was set to 250 µm.

#### Visualization and differential-expression

Gene expression was visualized with dot and violin plots. Dot plots displayed predefined gene sets across cell populations, with color showing scaled Z-scores (up-regulation only) and size reflecting expression frequency^109^. A continuous color gradient was applied to capture the full range of positive expression. Violin plots, generated with scCustomize^110^, were used to illustrate the expression distributions within each population.

For Visium HD, differential expression was computed with PrepSCTFindMarkers, and for the HSCA with FindMarkers, using ident.1 as the target cell type.

#### Cell type deconvolution of Visium HD data

To project single-cell resolved cell types from the HSCA core onto the Visium HD dataset, we applied reference-based deconvolution using RCTD^38^. The HSCA core object from Figure 5a served as reference, and the Visium HD dataset focused on the lower hair follicle region from donor D2. The RCTD reference object was constructed from raw UMI counts with cell type annotations, while the query object was built from Visium HD spot counts and spatial coordinates.

Deconvolution was run with doublet_mode = “doublet” to account for mixed-cell spots^38^. Posterior weights were stored as a predictions assay in the Visium HD Seurat object. For each spot, prediction scores across shared cell types were retained. To summarize contributions across the tissue, scores were aggregated by cell type and visualized as min-max normalized heatmaps using pheatmap^111^ with viridis scaling.

## CODE AVAILABILITY

The code used for the construction of the HSCA project is available at https://github.com/TolgaDuz/HSCA. The datasets are available on Zenodo (see Data Availability) and will further be made available for interactive exploration via the CellxGene browser. Updated links and implementations will be continuously provided on the GitHub repository.

## DATA AVAILABILITY

The HSCA provides a collection of objects (HSCA core, HSCA extended, pilosebaceous unit, lower hair follicle, and sebaceous gland objects) that contain raw and normalized counts, integrated embeddings, cell type annotations, and metadata (see Supplementary Table 4 for a detailed description of all data slots). These objects are available both as .rds files for R users and as equivalent AnnData objects for

Python users, allowing users to work in their preferred environment. All objects can be downloaded via Zenodo: https://doi.org/10.5281/zenodo.17088022.

Information on the original, published datasets included in the HSCA, necessary to access them, is provided in Supplementary Table 1.

## CONFLICT OF INTEREST

T.D., S.M.E.M., B.A., S.G. and N.H. are employees of Beiersdorf AG. D.T. and J.B. received consultation fees from Beiersdorf AG. The other authors declare no competing interests.

## ACKNOWLEDGEMENTS

This work was funded by Beiersdorf AG. D.T. was supported by the Hungarian National Research, Development and Innovation Office FK-132296 and ANN 139589.

## AUTHOR CONTRIBUTIONS

T.D. conceived and designed the study, with input from N.H. T.D. built the atlas, performed all analyses, generated the visualizations, and wrote the manuscript. S.M.E.M. and T.D. implemented and conducted the scIB pipeline. N.H. and J.B. supervised the project. N.H., J.B., M.O. and D.T. provided critical feedback and contributed to discussions. D.T. additionally contributed clinical expertise. B.A. and S.G. provided funding.

---

**Supplementary Fig. 1:**
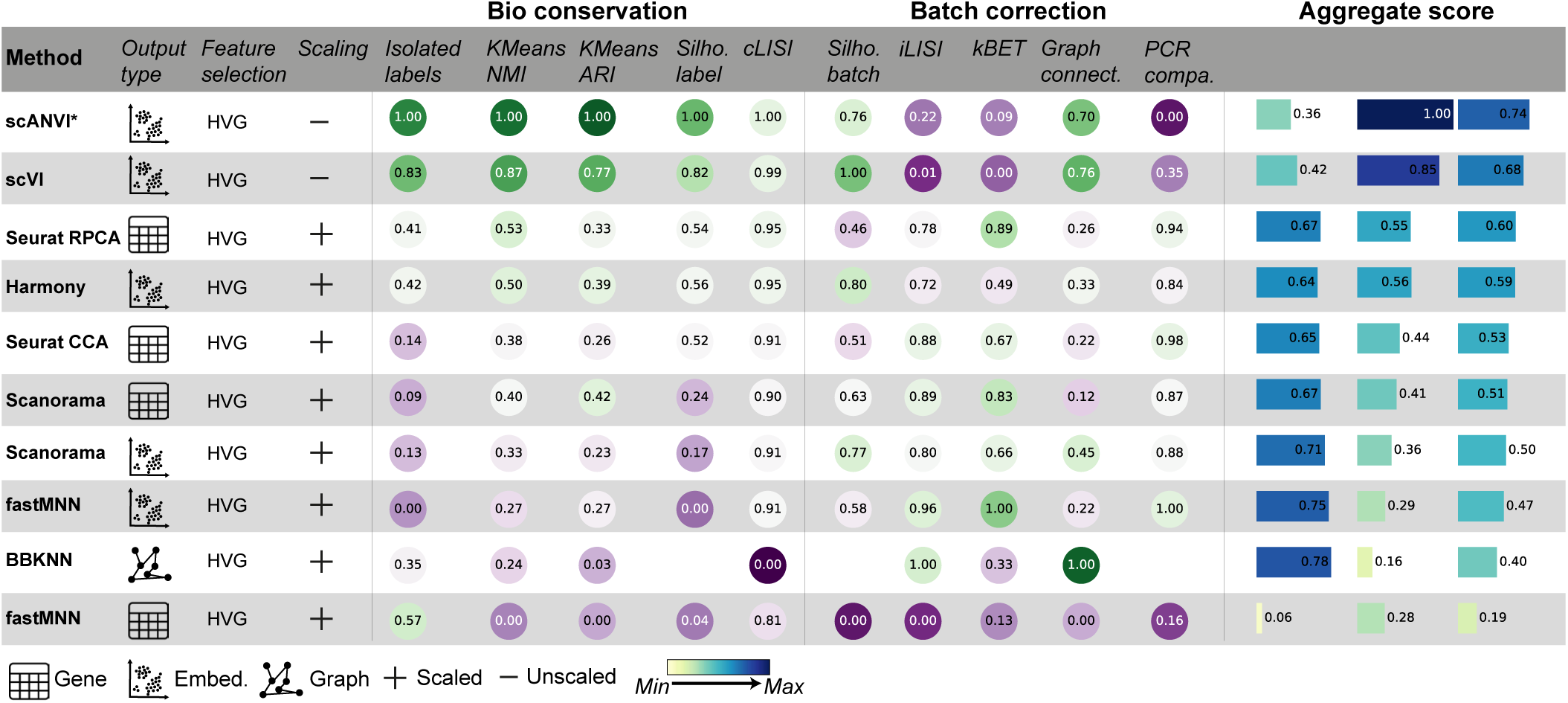
Benchmarking of integration methods with scIB for the HSCA core datasets15,22-32. For each method, 2000 highly variable genes were selected. The figure reports whether gene values were scaled before integration and specifies the type of output generated by the method, i.e. corrected expression values, a joint low-dimensional embedding, or an integrated graph. Certain tools such as Scanorama and fastMNN provide multiple output formats and were therefore evaluated under both conditions. Integration methods are ranked by their overall performance, which combines measures of batch correction and biological signal preservation into a single weighted score. In the case of scANVI, coarse cell type annotations were provided to guide the integration.

**Supplementary Fig. 2:**
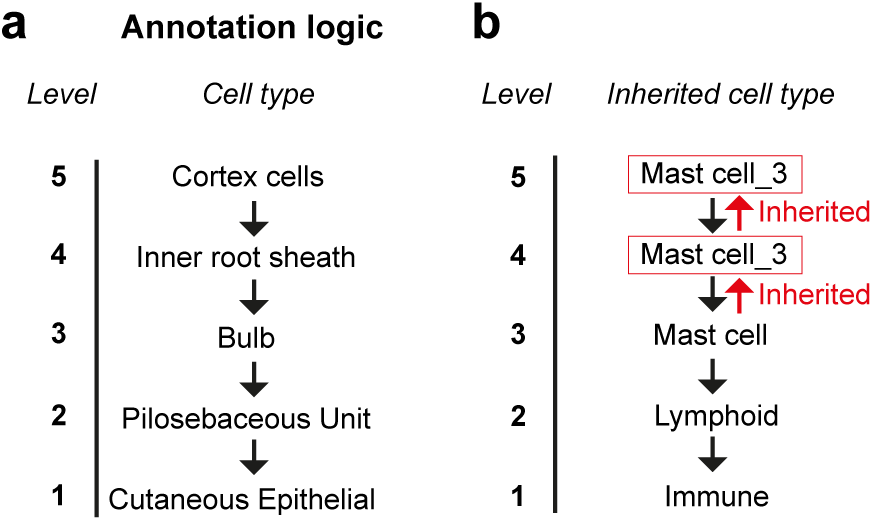
Bottom-up hierarchical cell type annotation in the HSCA. (**a**) Example of the finest annotation level (level 5) for cortex cells, with successive label propagation to inner root sheath (level 4), bulb (level 3), PSU (level 2), and cutaneous epithelial compartments (level 1), ensuring hierarchical consistency. (**b**) Illustration of the “inherited cell type” strategy, inspired by hierarchical annotation schemes established in the Lung Cell Atlas (Sikkema et al., 2023), and adapted here for skin. Cells whose most granular identity is defined at an intermediate level (e.g., mast cells) are propagated to lower hierarchical levels, with a numeric suffix indicating the original classification level.

**Supplementary Fig. 3:**
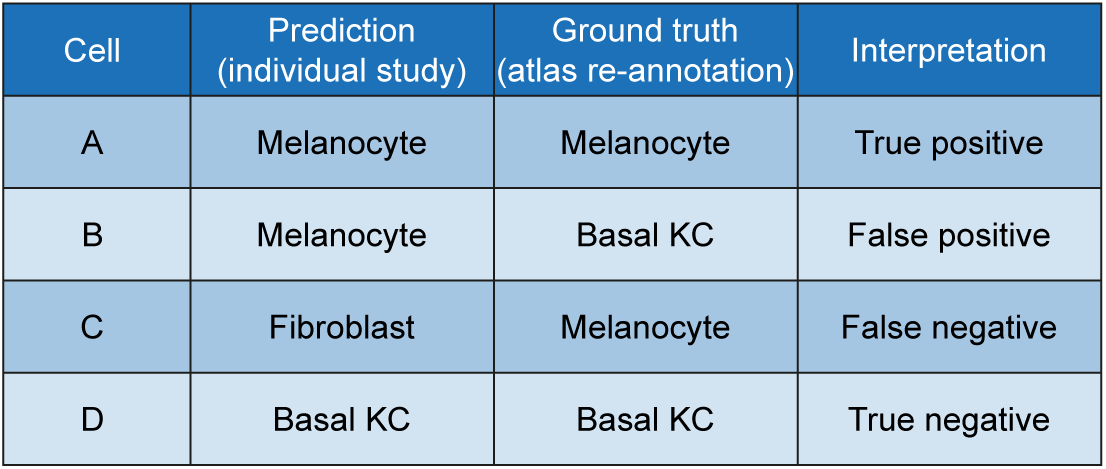
Confusion matrix of dataset-level versus atlas annotations. Comparison of dataset-specific annotations (treated as predictions) versus atlas-derived annotations from the HSCA core (treated as ground truth). True positives indicate cells correctly labeled in both the original dataset and the atlas (e.g., melanocytes annotated as melanocytes), false positives represent cells labeled as a given type in the original dataset but classified differently in the atlas, and false negatives are cells assigned to a given type in the atlas but differently in the original dataset. True negatives are not relevant in this context.

**Supplementary Fig. 4:**
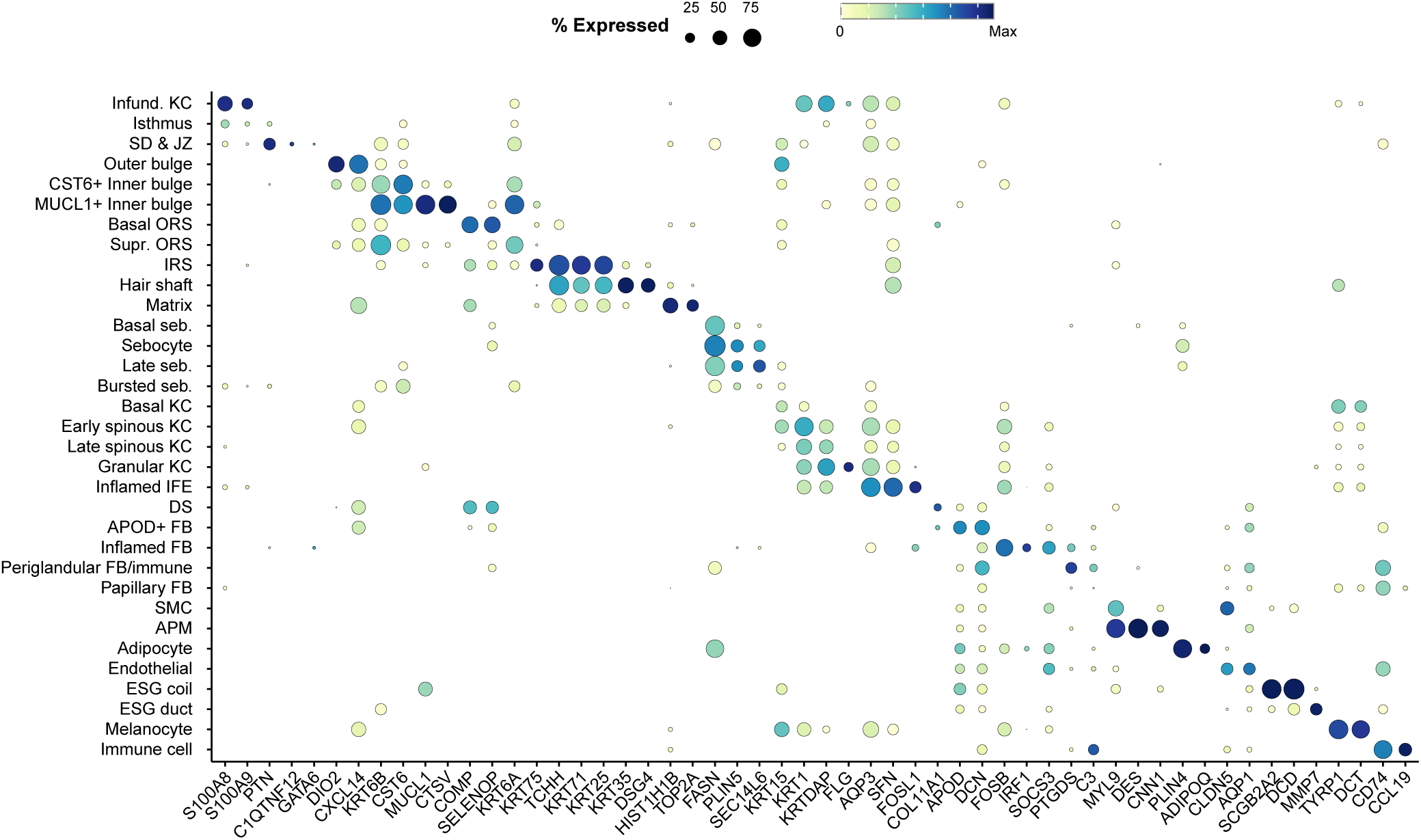
Marker gene expression across cell types in Visium HD. Dot plot showing expression of marker genes for distinct cell types at a binning resolution of 8 µm (corresponding to Figure 3). Abbreviations: see Supplementary Table 3.

**Supplementary Fig. 5:**
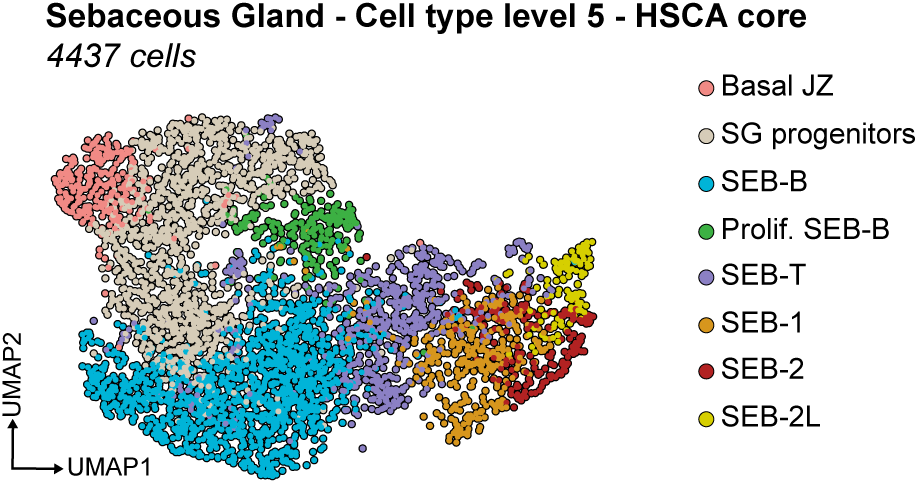
UMAP embedding of sebaceous gland differentiation stages, shown for comparison to PHATE (Fig. 5e).

**Supplementary Fig. 6:**
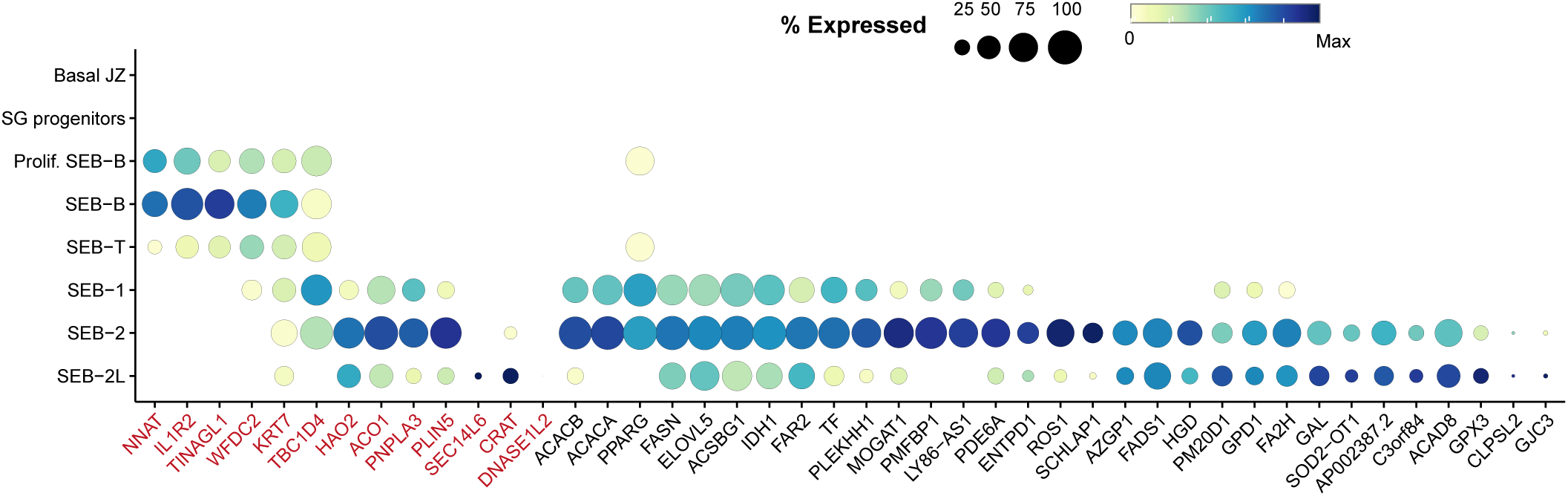
Dot plot of marker gene expression across sebocyte maturation stages. Dot plot displaying expression of marker genes associated with sebaceous gland differentiation stages resolved in the HSCA core. Genes highlighted in red correspond to stage-specific markers previously reported by (Düz et al., 2025), while additional markers identified in the HSCA core further refine the characterization of late-stage sebocytes.

**Supplementary Fig. 7:**
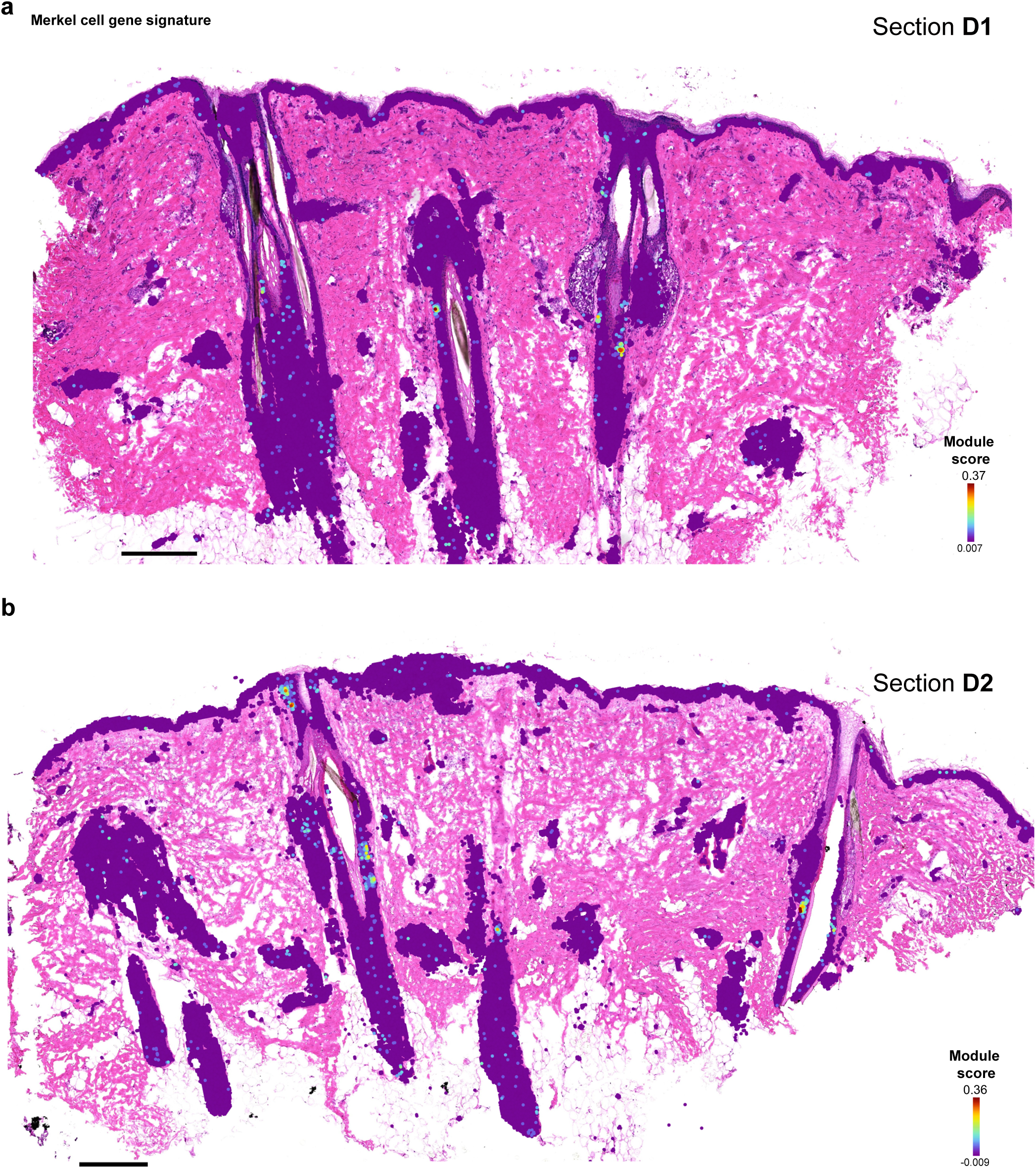
Visualization of the gene signature from Fig. 6b using the AddModuleScore function in Seurat. The signature was calculated excluding PCP4 and GNG4, as these genes displayed non-specific expression in eccrine sweat glands and muscle cells. Panel a shows tissue section D1, and panel b shows tissue section D2.

**Supplementary Fig. 8:**
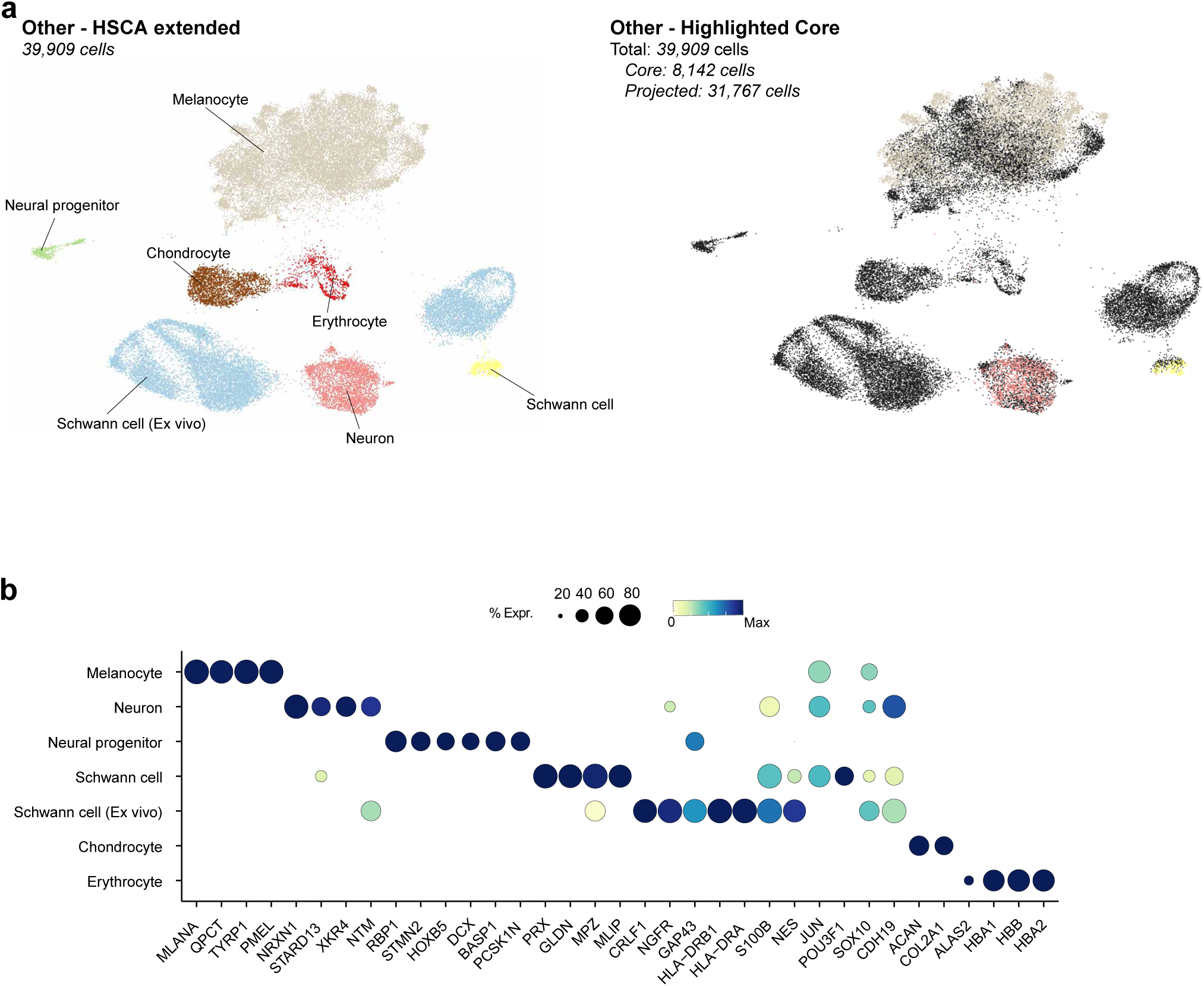
Other non-epidermal lineages in the HSCA extended. (**a**) UMAP of other lineages from the HSCA extended, including melanocytes, neurons, neural progenitors, Schwann cells, ex vivo Schwann cells, chondrocytes, and erythrocytes. Left: annotation of these subtypes. Right: HSCA core (colored) overlaid on the extended dataset (black). The extended dataset enables the identification of rare or previously absent cell types. (**b**) Dot plot of canonical marker genes used to annotate the cell types shown in panel a. Abbreviations: see Supplementary Table 3.\

**Supplementary Fig. 9:**
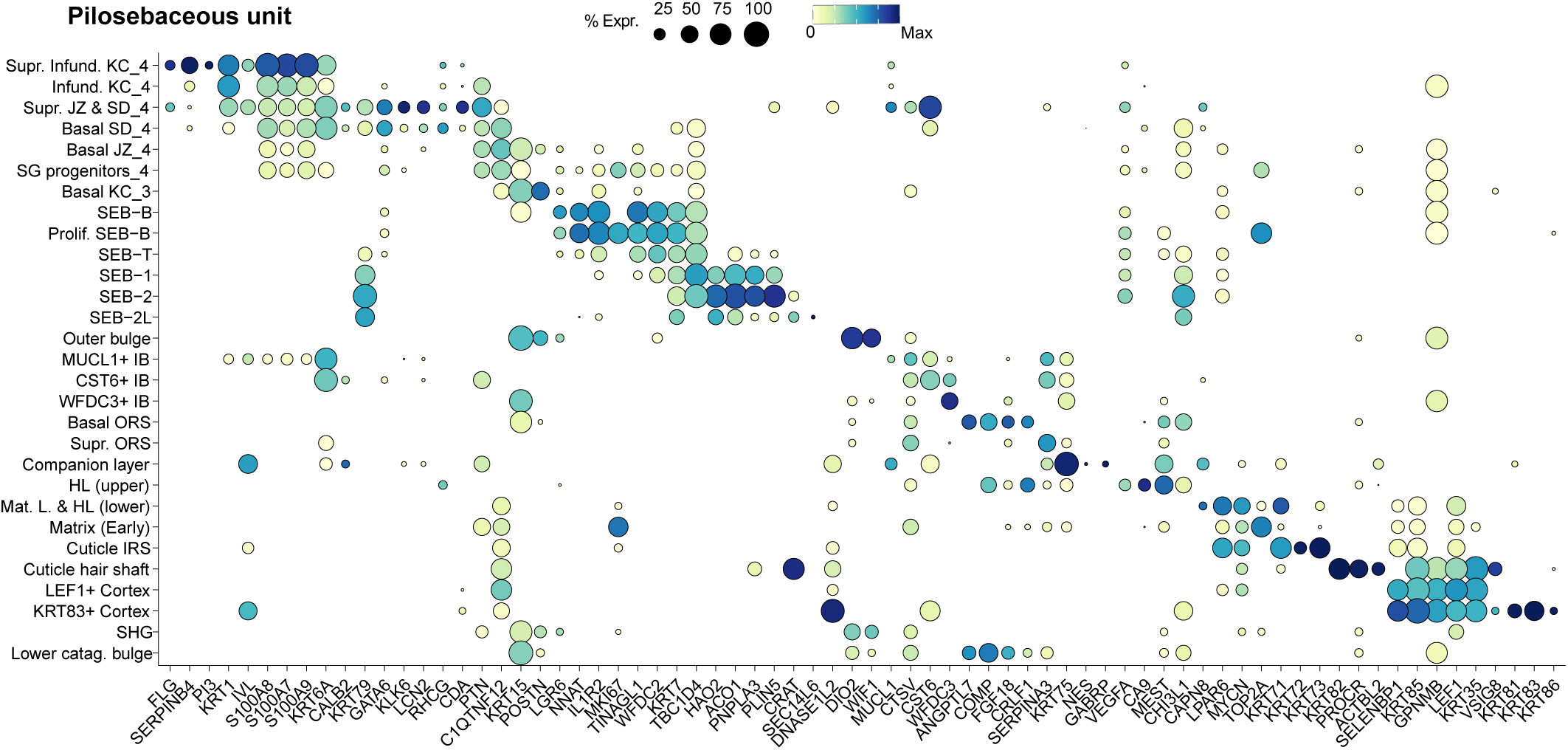
Dot plot of canonical marker genes used to annotate the cell types from the PSU. Abbreviations: see Supplementary Table 3.

**Supplementary Fig. 10:**
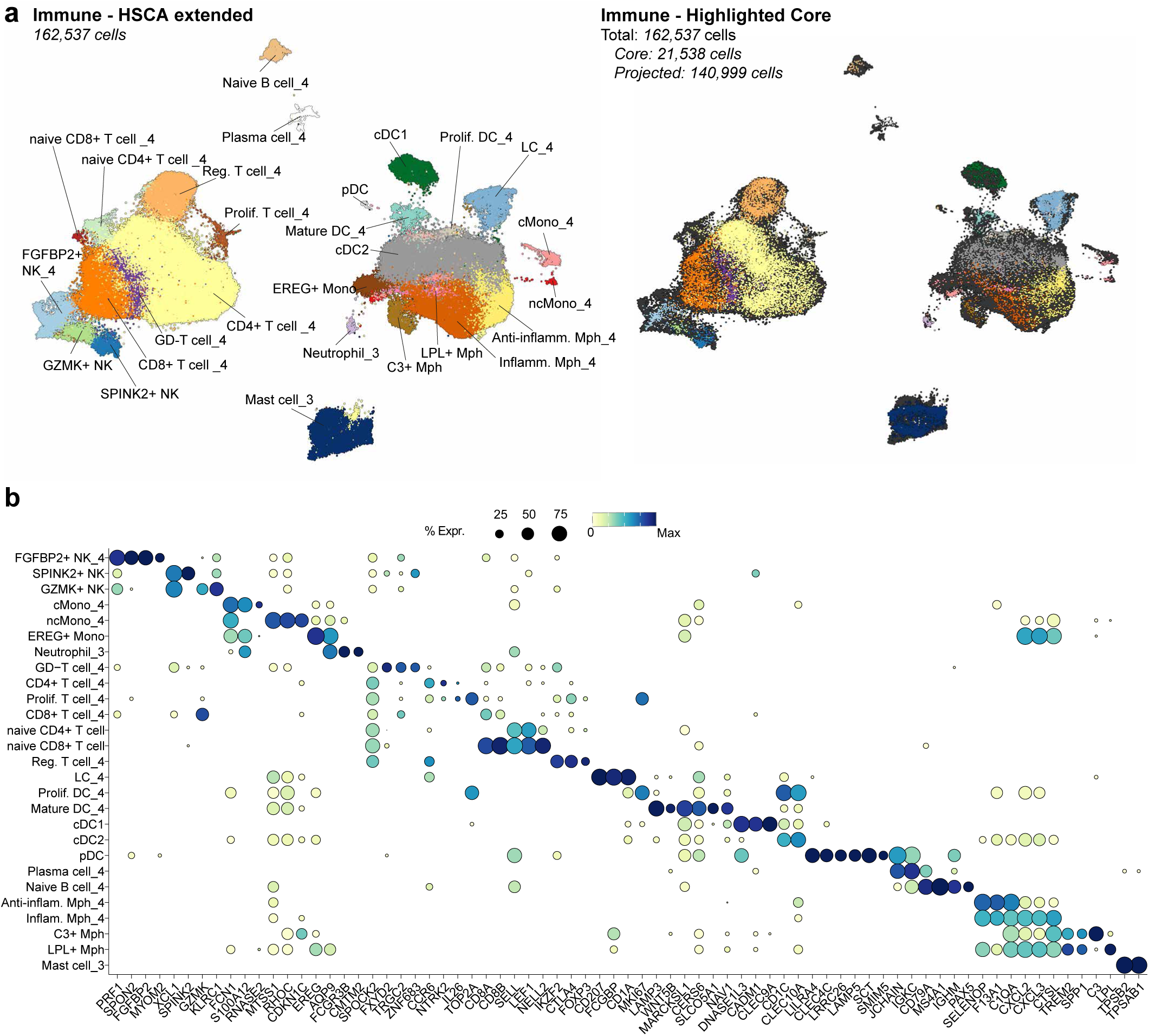
Immune cell types in the HSCA extended. (**a**) UMAP of immune cells from the HSCA extended. Left: annotation of immune inherited cell subtypes at level 5. Right: highlighting the HSCA core (colored) overlaid on the extended dataset (black), which illustrates strong concordance of cluster assignments while also revealing additional biological variance. The extended dataset enables the identification of novel cell types absent from the core, including naïve CD8+ and CD4+ T cells, a finer subdivision of naïve B cells, plasmacytoid dendritic cells, and increased heterogeneity within classical monocytes. (**b**) Dot plot of canonical marker genes used for annotation of the immune cell subtypes shown in panel a. Abbreviations: see Supplementary Table 3.

**Supplementary Fig. 11:**
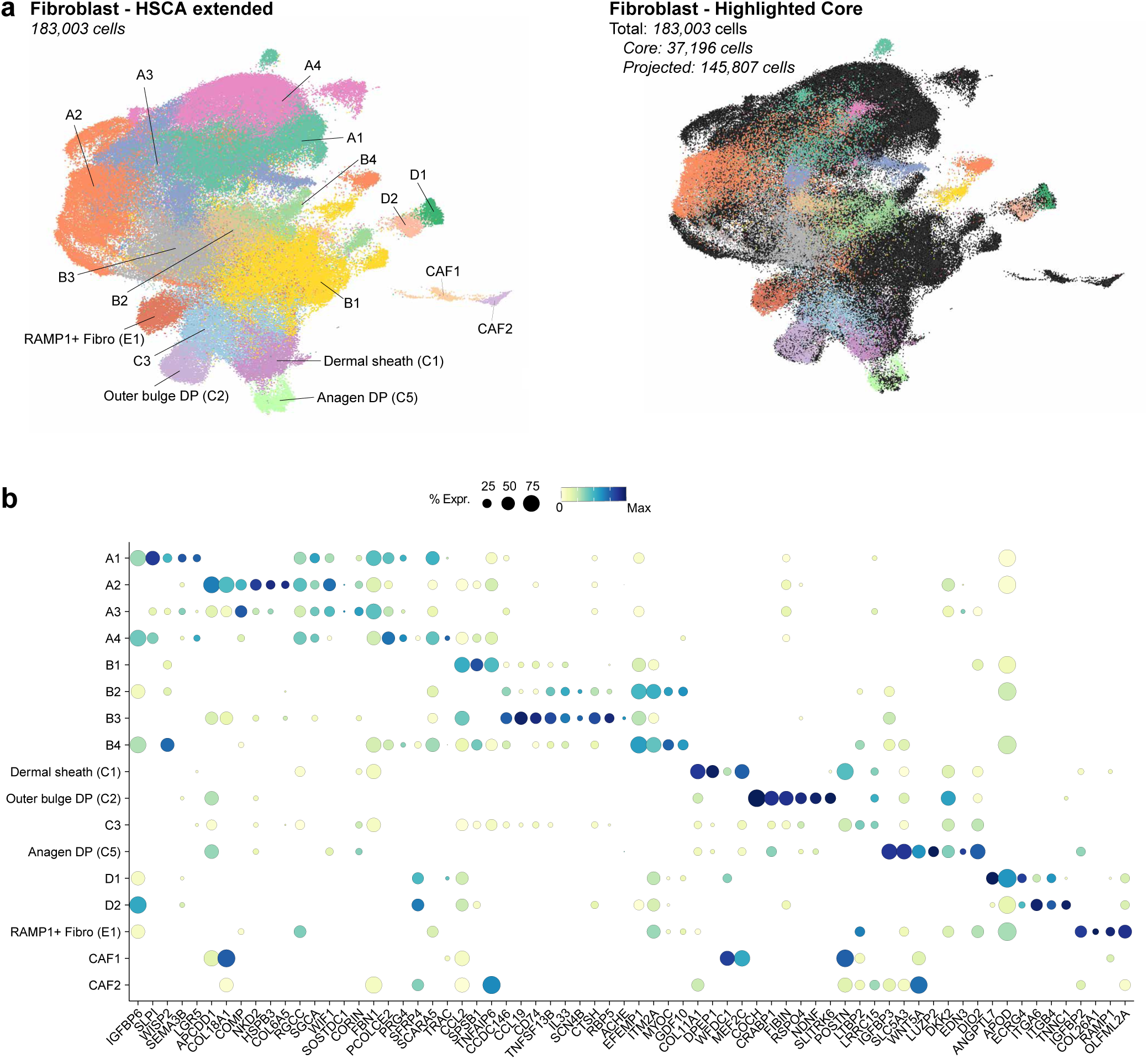
Fibroblast cell types in the HSCA extended. (**a**) UMAP of fibroblasts from the HSCA extended. Left: annotation of fibroblast subtypes at level 4. Right: HSCA core (colored) overlaid on the extended dataset (black), illustrating strong concordance of cluster assignments while also revealing additional biological variance. The extended dataset enables the identification of CAFs. (**b**) Dot plot of canonical marker genes used to annotate the fibroblast subtypes shown in panel a. Abbreviations: see Supplementary Table 3.

**Supplementary Fig. 12:**
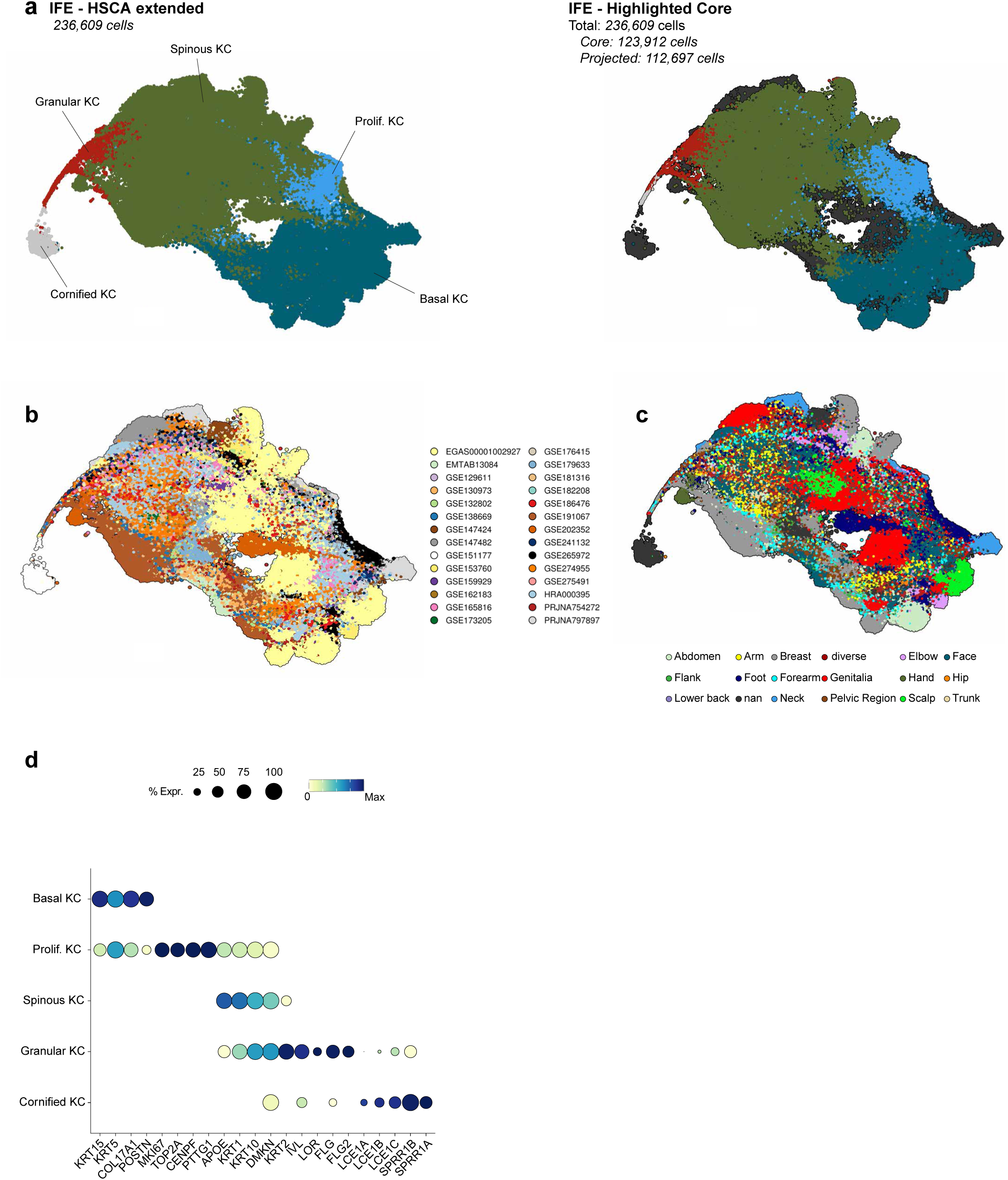
IFE cell types in the HSCA extended. (**a**) UMAP of IFE cells from the HSCA extended. Left: annotation of inherited IFE subtypes at level 3. Right: projection of the HSCA core (colored) onto the extended dataset (black), demonstrating strong concordance of cluster assignments while also capturing additional biological variability. The extended dataset further enables identification of a cornified keratinocyte cluster, observed specifically in dataset GSE151177, which was generated using an emigration protocol rather than enzymatic digestion. (**b**) UMAP of IFE cells from the HSCA extended, grouped by accession source. (**c**) UMAP of IFE cells grouped by anatomical region (level 2), highlighting the contribution of skin location to epidermal diversity. (**d**) Dot plot of canonical marker genes used to annotate IFE subtypes shown in panel (a). Abbreviations are listed in Supplementary Table 3.

**Supplementary Fig. 13:**
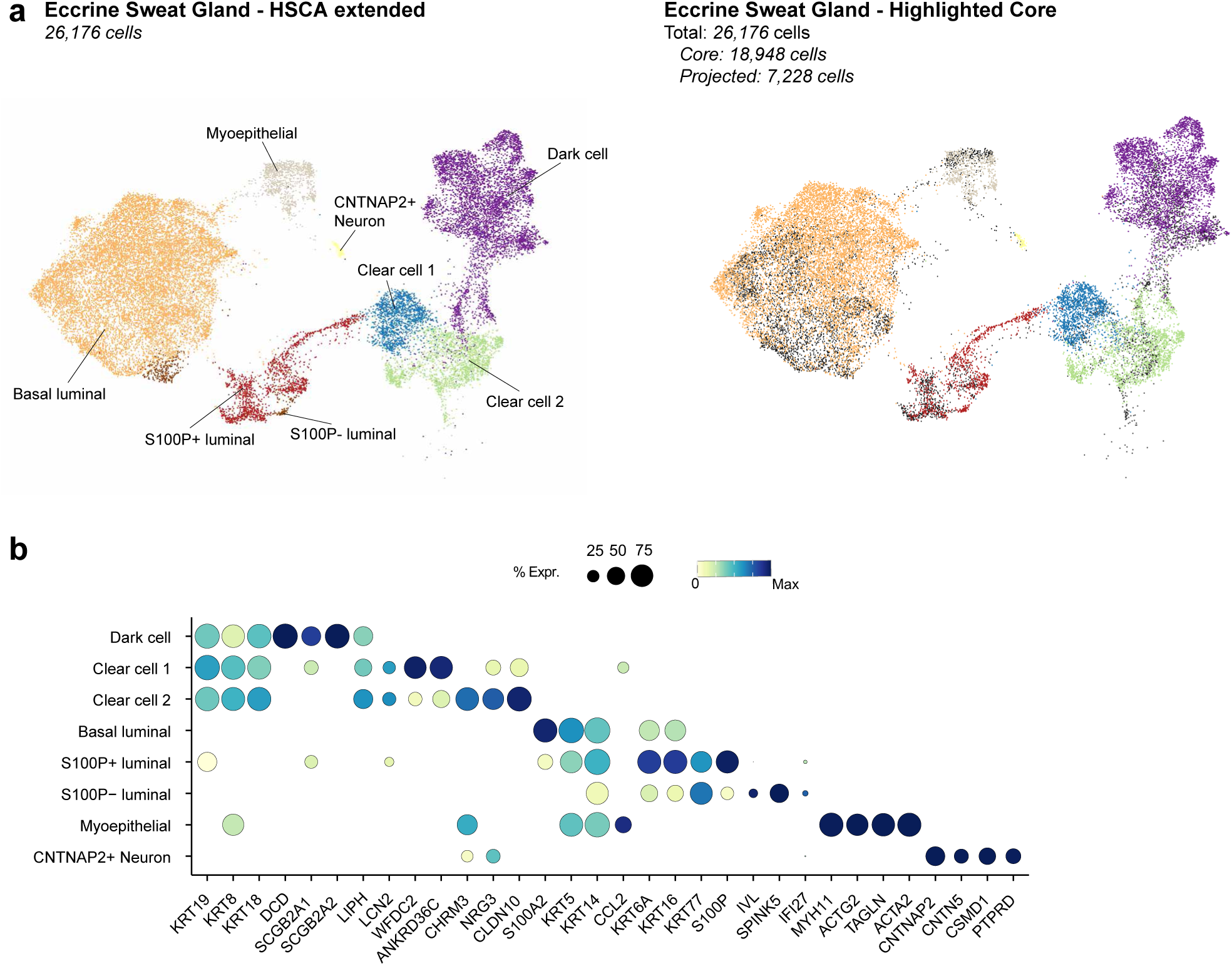
Eccrine sweat gland cell types in the HSCA extended. (**a**) UMAP of eccrine sweat gland cells from the HSCA extended. Left: annotation of eccrine sweat gland subtypes at level 4. Right: HSCA core (colored) overlaid on the extended dataset (black), illustrating strong concordance of cluster assignments while also revealing additional biological variance. (**b**) Dot plot of canonical marker genes used to annotate the eccrine sweat gland subtypes shown in panel a. Abbreviations: see Supplementary Table 3.

**Supplementary Fig. 14:**
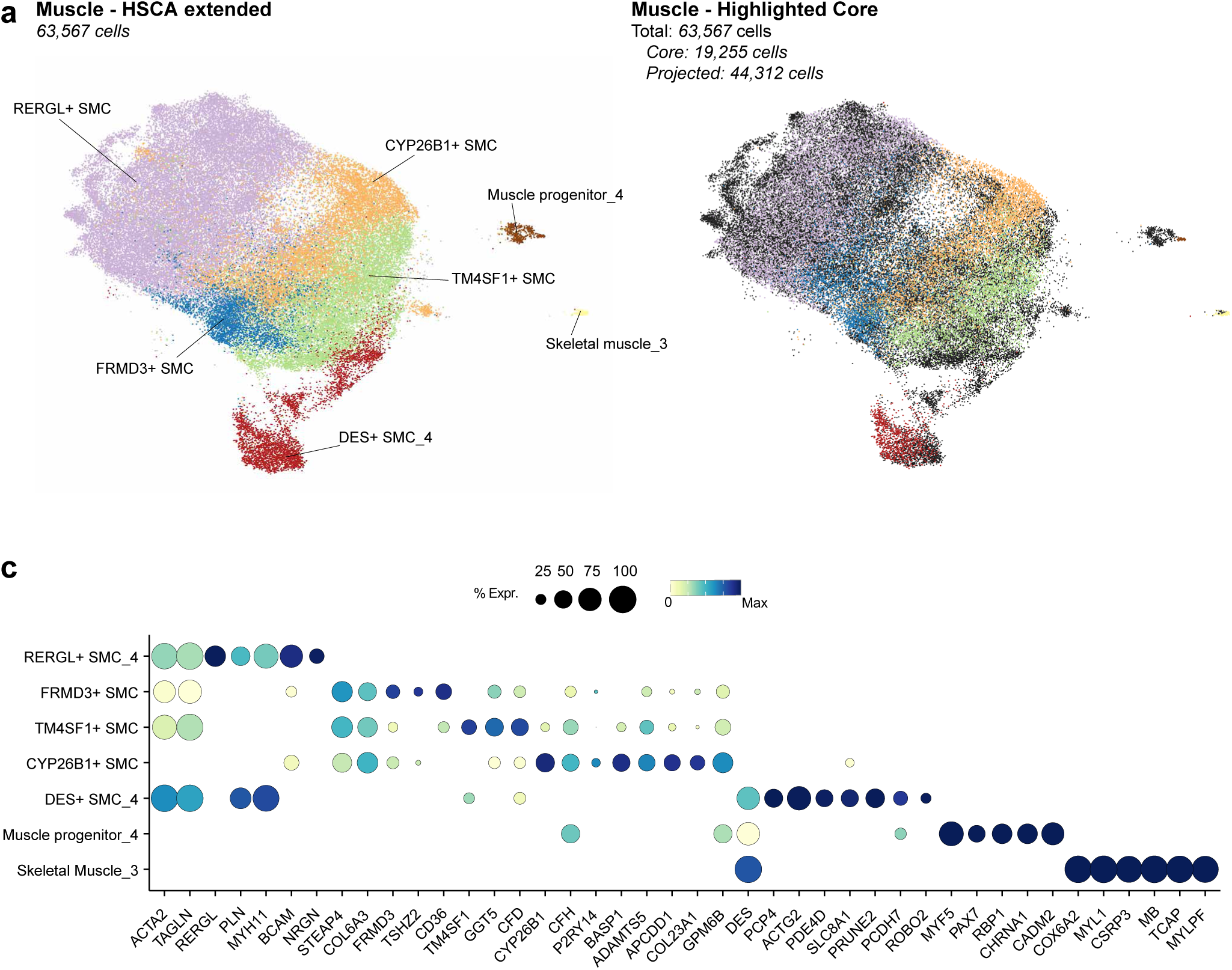
Muscle cell types in the HSCA extended. (**a**) UMAP of muscle cells from the HSCA extended. Left: annotation of muscle inherited cell subtypes at level 5. Right: HSCA core (colored) overlaid on the extended dataset (black), illustrating strong concordance of cluster assignments while also revealing additional biological variance. The extended dataset enables the identification of progenitor muscle cells. (**b**) Dot plot of canonical marker genes used to annotate the muscle cell subtypes shown in panel a. Abbreviations: see Supplementary Table 3.

**Supplementary Fig. 15:**
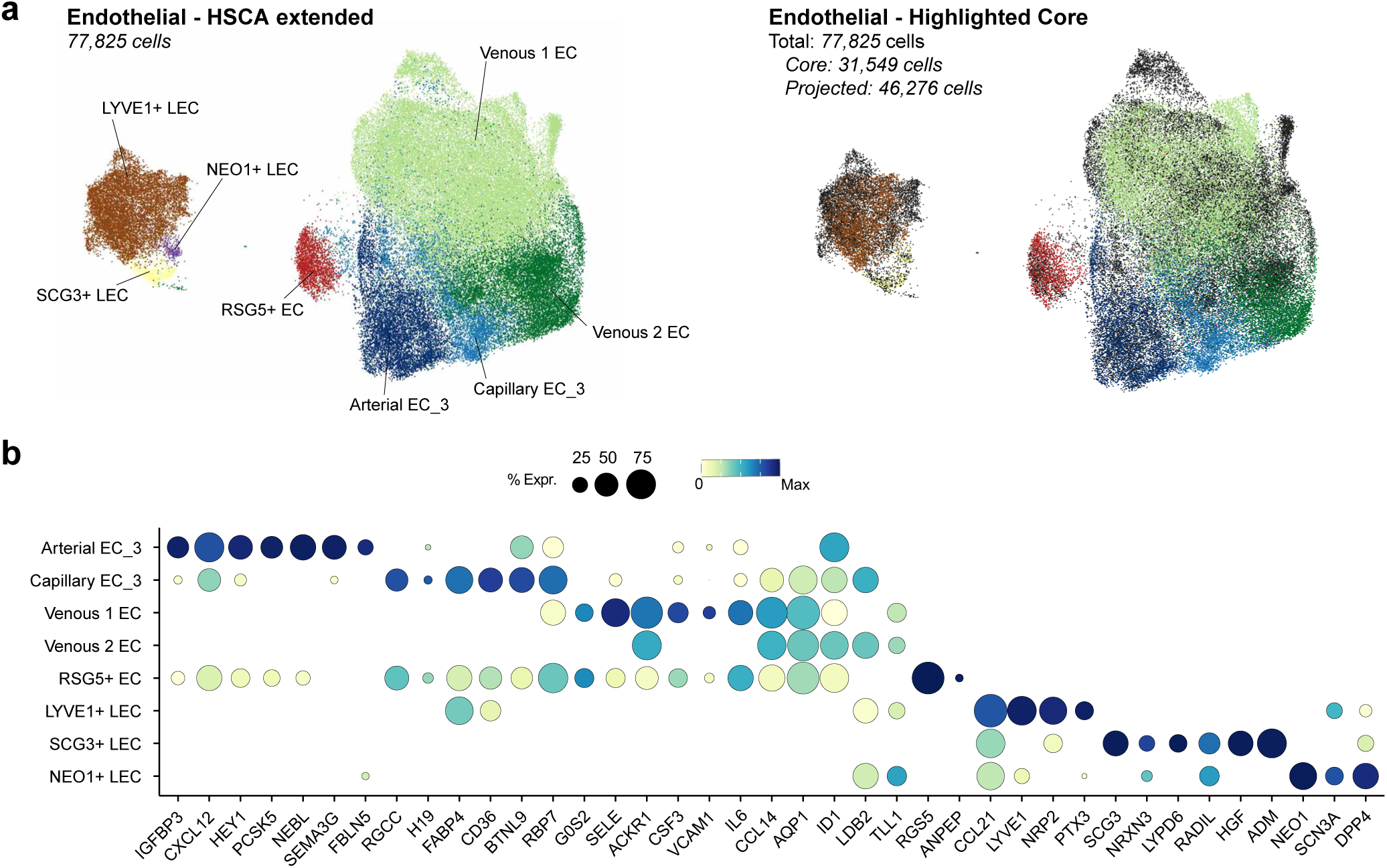
Endothelial cell types in the HSCA extended. (**a**) UMAP of endothelial subtypes from the HSCA extended. Left: annotation of endothelial subtypes at level 4. Right: HSCA core (colored) overlaid on the extended dataset (black), illustrating strong concordance of cluster assignments while also revealing additional biological variance. (**b**) Dot plot of canonical marker genes used to annotate the endothelial subtypes shown in panel a. Abbreviations: see Supplementary Table 3.

**Supplementary Fig. 16:**
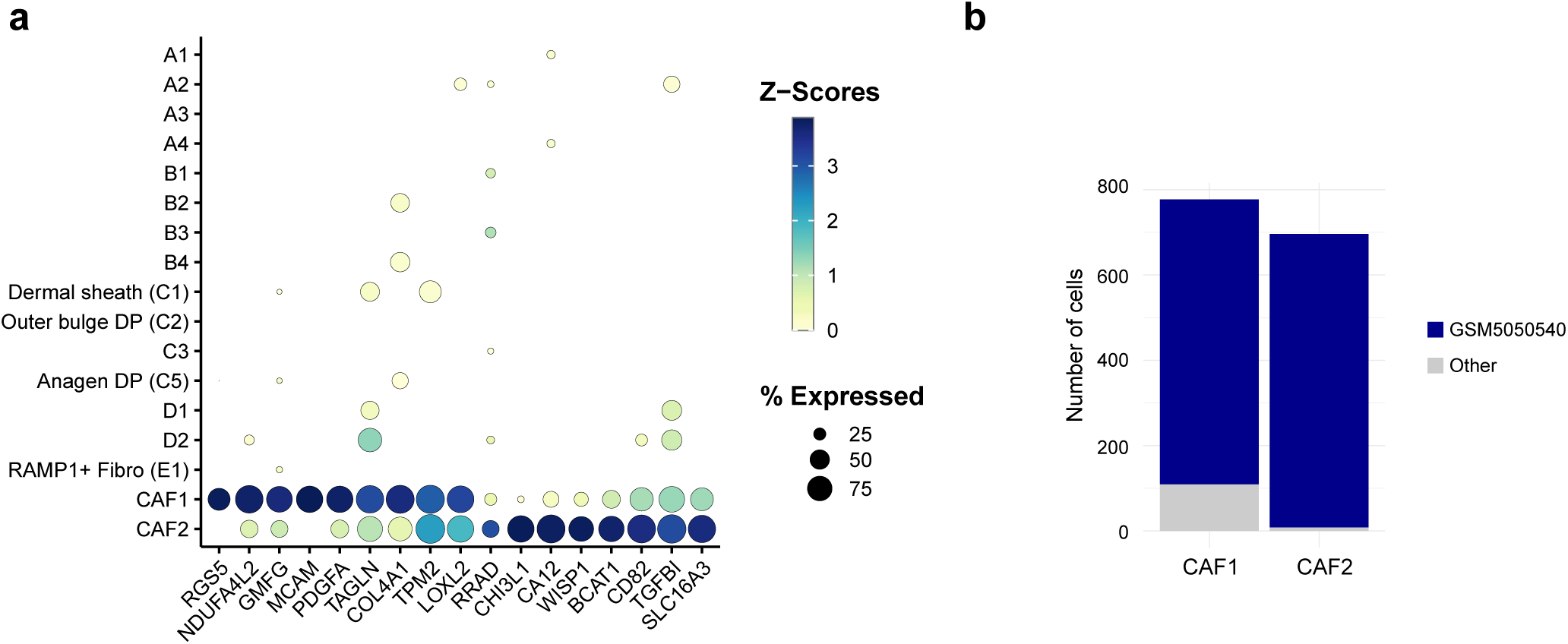
CAF signature and sample distribution in CAF clusters. (**a**) Dot plot showing canonical marker genes for cancer-associated fibroblasts (CAFs). (**b**) Bar plot showing sample composition of CAF clusters CAF1 and CAF2, highlighting that both clusters are dominated by a single sample (GSM5050540).

## Notes

https://doi.org/10.5281/zenodo.17088022

https://github.com/TolgaDuz/HSCA

